# Optogenetics and electron tomography for structure-function analysis of cochlear ribbon synapses

**DOI:** 10.1101/2022.05.10.491334

**Authors:** Rituparna Chakrabarti, Lina Maria Jaime Tobon, Loujin Slitin, Magdalena Redondo-Canales, Gerhard Hoch, Marina Slashcheva, Elisabeth Fritsch, Kai Bodensiek, Özge Demet Özçete, Mehmet Gültas, Susann Michanski, Felipe Opazo, Jakob Neef, Tina Pangrsic, Tobias Moser, Carolin Wichmann

## Abstract

Ribbon synapses of cochlear inner hair cells (IHCs) are specialized to indefatigably transmit sound information at high rates. To understand the underlying mechanisms, structure-function analysis of the active zone (AZ) of these synapses is essential. Previous electron microscopy studies of synaptic vesicle (SV) dynamics at the IHC AZ used potassium stimulation, which limited the temporal resolution to minutes. Here, we established optogenetic IHC stimulation followed by quick freezing within milliseconds and electron tomography to study the ultrastructure of functional synapse states with good temporal resolution. We characterized optogenetic IHC stimulation by patch-clamp recordings from IHCs and postsynaptic boutons revealing robust IHC depolarization and transmitter release. Ultrastructurally, the number of docked SVs increased and distances to the presynaptic density decreased upon short (17-25 ms) and long (48-76 ms) light stimulation paradigms. We did not observe enlarged SVs or other morphological correlates of homotypic fusion events. Our results suggest a rapid replenishment of docked SVs at IHC ribbon synapses and argue against synchronized multiquantal release under our experimental conditions.

## Introduction

Ribbon synapses of cochlear inner hair cells (IHCs) are specialized to maintain high release rates over prolonged periods of time. Their landmark structure, the synaptic ribbon, tethers several dozens of synaptic vesicles (SVs) and keeps them close to the active zone (AZ) membrane (Moser et al., 2019; Rutherford et al., 2021; Safieddine et al., 2012; Wichmann and Moser, 2015). Deciphering the mechanisms of SV release and replenishment in IHCs is required to understand their efficient and indefatigable glutamate release. Ultrastructural analysis of SV pools in defined functional states, such as during phasic or sustained transmitter release, is an important approach to investigate presynaptic mechanisms in general.

Numerous studies based on electron tomography (ET) describe the presence of morphologically docked SVs at central synapses (e.g. Hintze et al., 2021; Imig et al., 2014; Imig et al., 2020; Kusick et al., 2020; Maus et al., 2020; Siksou et al., 2007). At such conventional synapses, docked SVs are thought to constitute the readily-releasable pool (RRP) (Schikorski and Stevens, 1997), while SVs tethered to the AZ might represent morphological correlates for SV recruitment to the release sites (Cole et al., 2016; Fernández-Busnadiego et al., 2010; Fernández-Busnadiego et al., 2013; Maus et al., 2020; Siksou et al., 2007). Recruitment appears to involve a first step mediated by long tethers of up to 45 nm, followed by formation of shorter tethers which might correspond to the soluble N-ethylmaleimide-sensitive-factor attachment receptor (SNARE) complex (Cole et al., 2016; Fernández-Busnadiego et al., 2010; Imig et al., 2014). Therefore, morphological features like tethering or docking might reflect different functional states pf SVs *en route* to fusion.

Correlating function and structure ideally employs rapid immobilization of the synapses in defined functional states. Recently, SV dynamics were investigated by combining optogenetics with immobilization within milliseconds by high-pressure freezing (Opto- HPF). Optogenetics grants short, precise stimulation of neurons expressing the light-sensitive ion channel channelrhodopsin (ChR) (Nagel et al., 2002; Nagel et al., 2003). Such precise stimulation allowed to ultrastructurally resolve transient events of exo/endocytosis at several conventional synapses such as *C. elegans* neuromuscular junctions (Kittelmann et al., 2013; Watanabe et al., 2013a) and murine hippocampal synapses (Borges-Merjane et al., 2020; Imig et al., 2020; Watanabe et al., 2013b).

Until now, structure-function analysis of hair cell ribbon synapses relied on seconds to minutes range depolarization by high K^+^ (Chakrabarti et al., 2018; Jung et al., 2015b; Lenzi et al., 2002; Pangrsic et al., 2010; Strenzke et al., 2016). SVs situated close to the AZ membrane — referred to as the membrane-proximal (MP)-SV pool — are thought to represent the RRP, while SVs around the ribbon — ribbon-associated (RA)-SVs — seem to be recruited for release in a later phase (Lenzi et al., 2002). MP-SVs are often connected to the AZ membrane by tethers (Chakrabarti et al., 2018; Frank et al., 2010; Vogl et al., 2015) and seem to be organized in sub-pools based on the number and length of tethers. These sub-pools might represent different recruitment states of the SVs prior to docking. However, docked SVs are rare in IHCs at rest but become more frequent upon prolonged K^+^ depolarization (Chakrabarti et al., 2018). Yet, K^+^ depolarization does not enable a time-resolved analysis of exocytosis at IHC ribbon synapses.

Time-resolved analysis is also relevant when addressing the long-standing quest on whether SVs fuse in a coordinated manner or independently from each other in IHCs. From postsynaptic recordings of the spiral ganglion neurons, the high variability in the amplitude and shape of spontaneous excitatory postsynaptic currents (sEPSCs) was initially interpreted as the release of multiple SVs in a more or less synchronized manner (Glowatzki and Fuchs, 2002). The alternative, classical model of uniquantal release, was then proposed based on experiments and modeling (Chapochnikov et al., 2014), and further corroborated by direct measurements of single fusion events (Grabner and Moser, 2018). In the uniquantal release framework, amplitude and shape heterogeneity were attributed to glutamate release via a fusion pore with different progress towards full collapse fusion (Chapochnikov et al., 2014). Such different scenarios might be mirrored in the number of docked SVs, SVs sizes and the distribution of SV pools. For instance, coordinated multivesicular release by compound and/or cumulative fusion is expected to result in larger vesicles at the AZ.

Here, we implemented Opto-HPF of IHC ribbon synapses to capture the structural correlates of exocytosis. We modified a conventional high-pressure freezing machine (HPM) to control optical stimulation in correlation to freezing on a millisecond time scale. Our study revealed that upon depolarization (i) the number of docked SVs increases, (ii) MP-SVs reside closer to the AZ membrane, (iii) correlates of compound and/or cumulative fusion are lacking and (iii) the total number of RA-SVs remains unchanged. Our results constitute the first report of morphological correlates to exocytosis occurring within milliseconds of stimulation at this highly specialized synapse.

## Materials and Methods

### Animals

The mice were bred at the animal facility in the University Medical Center Göttingen (UMG). Animal handling and all experimental procedures were in accordance with the national animal care guidelines issued by the animal welfare committees of the University of Göttingen and the Animal Welfare Office of the State of Lower Saxony (AZ 509.42502/01-27.03).

For expression of ChR2-H134R-EYFP (Nagel et al., 2003) in IHCs, we crossbred the Ai32 mouse line (Madisen et al., 2012; RRID:IMSR_JAX:024109) with two different Vglut3-Cre mouse lines. The ChR2-H134R-EYFP construct is preceded by a STOP codon flanked by loxP sequences such that expression only commences upon Cre recombination (*cre^+^/cre^+^* or *cre^+^/cre^-^;* abbreviated *cre^+^*). The first ChR2 Vglut3-driven line, termed Ai32VC, used a previously published transgenic Vglut3-Cre line (Jung et al., 2015b). In the Ai32VC line, animals expressing ChR2 were either *fl/fl cre^+^* or *fl/+ cre^+^*, which we will abbreviate *Ai32VC cre^+^*. For the second ChR2 Vglut3-driven line, termed Ai32KI, we used Vglut3-Ires-Cre-KI mice (Lou et al., 2013; Vogl et al., 2016). ChR2 expressing animals were either *fl/fl cre^+^/cre^-^* or *fl/+ cre^+^/cre*^-^, which we will abbreviate *Ai32KI cre^+^*. Littermate controls from both lines (*fl/fl +/+*) are nicknamed WT. C57Bl6/J mice, are abbreviated “B6J”.

For immunohistochemistry analysis, three age groups of Ai32KI mice were used. The first group (G1) includes 4-5 months-old mice *Ai32KI cre^+^*, *N_animals_* = 3, n = 170 cells; and WT, *N_animals_* = 2, n = 99 cells. The second group (G2) corresponds to 6-7 months-old mice *Ai32KI cre^+^*, *N_animals_* = 2, n = 116 cells; and WT, *N_animals_* = 2, n = 87 cells. The third group (G3) includes 9-12 months-old mice *Ai32KI cre^+^*, *N_animals_* = 2, n = 129 cells; and WT, *N_animals_* = 2, n = 49 cells. For patch-clamp recordings, Ai32VC or Ai32KI mice were used at postnatal days (P) 14-20 (i.e. after the onset of hearing): 7 animals were *Ai32VC cre^+^* (*fl/fl*), 5 animals were *Ai32VC cre^+^* (*fl/+*) and 3 mice were *Ai32KI cre^+^* (*fl/fl*). For pre-embedding immunogold, we used *Ai32KI cre^+^*. For Opto-HPF, *Ai32VC cre^+^*, *Ai32KI cre^+^* and B6J mice were used as controls at P14-20 at different stimulation durations (see Table 1 below).

**Table 1.** Genotypes, animal numbers as well as the ages of the animals used in the experiments.

| Experiment | genotype | N<sub>animals</sub> | n (cells/ribbon) | age |
| --- | --- | --- | --- | --- |
| Immunohistochemistry | *Ai32KI cre<sup>+</sup>* | 3 | 170 cells | Group 1: 4-5 month |
|  | *Ai32KI cre<sup>+</sup>* | 2 | 116 cells | Group 2: 6-7 month |
|  | *Ai32KI cre<sup>+</sup>* | 2 | 129 cells | Group 3: 9-12 month |
| Immunohistochemistry | WT | 2 | 99 cells | Group 1: 4-5 month |
|  | WT | 2 | 87 cells | Group 2: 6-7 month |
|  | WT | 2 | 49 cells | Group 3: 9-12 month |
| Pre-embedding immunogold | *Ai32KI cre<sup>+</sup>* | 1 |  | P17 |
| Patch-clamp | *Ai32VC cre<sup>+</sup> (fl/fl)* | 7 |  | P14-17 |
|  | *Ai32VC cre<sup>+</sup> (fl/+)* | 5 |  | P14-17 |
|  | *Ai32KI cre<sup>+</sup>* | 3 | 21 cells | P16-20 |
| Electron tomography MP SVs |  |  |  | P14-20 |
| ShortStim | *Ai32VC cre<sup>+</sup>* | 1 | 11 ribbons |  |
| LongStim | *Ai32VC cre<sup>+</sup>* | 2 | 15 ribbons |  |
| LongStim | B6J | 2 | 15 ribbons |  |
| LongStim | *Ai32KI cre<sup>+</sup>* | 2 | 11 ribbons |  |
| NoLight | *Ai32VC cre<sup>+</sup>* | 2 | 9 ribbons |  |
| NoLight | *Ai32KI cre<sup>+</sup>* | 2 | 8 ribbons |  |
| Electron tomography RA SVs |  |  |  | P14-20 |
| ShortStim | *Ai32VC cre<sup>+</sup>* | 1 | 11 ribbons |  |
| LongStim | *Ai32VC cre<sup>+</sup>* | 1 | 10 ribbons |  |
| LongStim | B6J | 1 | 9 ribbons |  |
| LongStim | *Ai32KI cre<sup>+</sup>* | 2 | 11 ribbons |  |
| NoLight | *Ai32VC cre<sup>+</sup>* | 2 | 9 ribbons |  |
| NoLight | *Ai32KI cre<sup>+</sup>* | 2 | 8 ribbons |  |

Some Ai32VC animals showed germline recombination of the EGFP cassette from the CGCT construct (coming from the Vglut3-Cre line) and/or germline recombination of the cassette ChR2-H134R-EYFP due to unspecific Cre-recombinase activity. This resulted in the ectopic expression (e.g. in non-IHCs in the cochlea) of EGFP and/or ChR2-H134R- EYFP in the absence of Cre-recombinase that could be observed using immunohistochemistry. Additional primers were designed to detect mice with general recombination during genotyping: Ai32 recombinant (forward primer 5’ – GTGCTGTC TCATCATTTTGGC – 3‘, and reverse primer 5’ – TCCATAATCCATGGTGGCAAG – 3‘) and CGCT recombinant (forward primer 5’ – CTGCTAACCATGTTCATGCC – 3‘, and reverse primer 5’ – TTCAGGGTCAGCTTGCCGTA – 3‘). The genotype of the animals was determined before the onset of hearing and further corroborated post-mortem.

### Patch-clamp recordings

Perforated patch-clamp recordings from IHCs expressing ChR2-H134R-EYFP were performed as described previously (Moser and Beutner, 2000). Briefly, the apical coils of the organ of Corti were dissected from euthanized mice at P14-20 in HEPES Hank’s solution containing 5.36 mM KCl, 141.7 mM NaCl, 1 mM MgCl_2_-6H_2_O, 0.5 mM MgSO_4_- 7H_2_O, 10 mM HEPES, 0.5 mg/ml L-glutamine, and 1 mg/ml D-glucose, pH 7.2, osmolarity ∼300 mOsm/l. By removing some supporting cells, the basolateral face of the IHCs was exposed and patch-clamp was established using Sylgard™-coated 1.5 mm borosilicate pipettes. The intracellular pipette solution contained: 135 mM KCl, 10 mM HEPES, 1 mM MgCl_2_, and 300 μg/ml amphotericin B (osmolarity ∼290 mOsm/l). The organ of Corti was bathed in an extracellular solution containing 126 mM NaCl, 20 mM TEA-Cl, 2.8 mM KCl, 2 mM CaCl_2_, 1 mM MgCl_2_, 1 mM CsCl, 10 mM HEPES, and 11.1 mM D-glucose, pH 7.2, osmolarity ∼300 mOsm/l. All patch-clamp recordings were performed at room temperature (20-25°C). An EPC-9 amplifier (HEKA electronics) controlled by Pulse or Patchmaster software (HEKA electronics) was used for the measurements. Currents were leak corrected using a p/10 protocol. IHCs with leak currents exceeding -50 pA at - 84 mV holding potential or with a series resistance higher than 30 MΩ were excluded from the analysis. A red filter was positioned between the light source and the recording chamber to avoid partial depolarization of the IHCs by the transillumination light.

In order to assess optogenetically evoked IHC exocytosis, postsynaptic recordings from afferent boutons were performed as described previously (Glowatzki and Fuchs, 2002; Huang and Moser, 2018). Whole-cell patch clamp recordings from the postsynaptic bouton was established using heat-polished, Sylgard™-coated 1 mm thin-glass borosilicate pipettes. The intracellular solution contained 137 mM KCl, 5 mM EGTA, 5 mM HEPES, 1 mM Na_2_-GTP, 2.5 mM Na_2_-ATP, 3.5 mM MgCl_2_·6H_2_O and 0.1 mM CaCl_2_, pH 7.2 and osmolarity of ∼290 mOsm/l. Boutons with leak currents exceeding -100 pA at -94 mV holding potential were excluded from the analysis. The series resistance of the bouton recordings was calculated offline as reported in (Huang and Moser, 2018). Recordings with a bouton series resistance > 80 MΩ were discarded.

### Photostimulation for cell-physiology

Photostimulation of IHCs was achieved using a blue 473 nm laser (MBL 473, CNI Optoelectronics). Irradiance and duration of the light pulses were controlled using the EPC-9 amplifier via a custom controller unit, allowing the transformation of a particular voltage to a particular laser power (i.e. photostimulation during 5, 10 or 50 ms from 2 to 5 V with different increasing steps). A FITC filter set was used to direct the stimulating blue light to the sample. Radiant flux (mW) was measured before each experiment with a laser power meter (LaserCheck; Coherent Inc. or Solo2 Gentec-eo) placed under the 40x objective lens. The diameter of the illumination spot was estimated using a green fluorescent slide and a stage micrometer, and it was used to calculate the irradiance in mW/mm^2^. Photocurrents were measured in voltage-clamp mode and the photodepolarization in current-clamp mode. Only the first series of evoked photocurrents and photodepolarizations were analyzed in order to rule out potential changes in the photoresponses due to inactivation of ChR2-H134R-EYFP (Lin et al., 2009).

### Immunohistochemistry

Freshly dissected apical turns of the organ of Corti (as described above) were fixed on ice for 60 min with 4% formaldehyde in Phosphate-Buffered Saline (PBS: 137 mM NaCl, 2.7 mM KCl, 8 mM N_2_HPO_4_, 0.2 mM KH_2_PO_4_), followed by 3x10 min wash with PBS. A blocking step was performed for 1 h at room temperature with goat serum dilution buffer (GSDB: 16% goat serum, 450 mM NaCl, 0.3% Triton X-100, and 20 mM phosphate buffer, pH 7.4). Afterwards, the samples were incubated overnight in a wet chamber at 4°C with the following GSDB-diluted primary antibodies: chicken anti-GFP (1:500, Abcam, ab13970; RRID:AB_300798), rabbit anti-myo6 (1:200, Proteus Biosciences, 25-6791; RRID:AB_10013626), mouse anti-CtBP2 (1:200, BD Biosciences, 612044; RRID:AB_399431), mouse anti-neurofilament 200 (1:400, Sigma, N5389; RRID:AB_260781) and rabbit anti-Vglut3 (1:300, SySy, 135 203; RRID:AB_887886). After 3x10 min wash with wash buffer (450 mM NaCl, 0.3% Triton X-100, and 20 mM phosphate buffer, pH 7.4), GSDB-diluted secondary antibodies were applied for 1 h at room temperature: goat anti-chicken Alexa Fluor 488 (1:200, Invitrogen, A11039; RRID:AB_2534096), AbberiorStar 580 goat conjugated anti-rabbit (1:200, Abberior, 2- 0012-005-8; RRID:AB_2810981), AbberiorStar 635p goat conjugated anti-mouse (1:200, Abberior, 2-0002-007-5; RRID:AB_2893232), goat anti-mouse Alexa Fluor 647 (1:200, Invitrogen, A-21236; RRID:AB_2535805) and goat anti-rabbit Alexa Fluor 568 (1:200, ThermoFisher, RRID:AB_143157). A final washing step was done for 3x10 min with wash buffer and, exclusively in Ai32VC samples, for 1x10 min in 5 mM phosphate buffer. The samples were mounted onto glass slides with a drop of mounting medium (Mowiol® 4- 88, Roth) and covered with glass coverslips.

Confocal images were acquired using an Abberior Instruments Expert Line STED microscope with a 1.4 NA 100x oil immersion objective and with excitation lasers at 488, 561, 594 and 633 nm. Images were processed using the FIJI software (Schindelin et al., 2012) and assembled with Adobe Illustrator Software.

### Immunogold pre-embedding

In order to verify the membrane localization of ChR2 within the IHC, we performed pre-embedding immunogold labeling using nanogold (1.4 nm gold)-coupled nanobodies (information see below) for the line *Ai32KI cre^+^*(*N* = 1) on a freshly dissected organ of Corti. The labeling was essentially done as described in (Strenzke et al., 2016) with a few modifications. Samples were fixed in 2% paraformaldehyde with 0.06% glutaraldehyde in 1x piperazine-N,N′-bis(2-ethanesulfonic acid)-EGTA-MgSO_4_ (PEM) solution (0.1 M PIPES; 2 mM EGTA; 1 mM MgSO_4_ x 7H_2_O) for 90 min on ice and subsequently washed twice for 15 min each in 1x PEM at RT. Next, the samples were blocked for 1 h in 2% BSA / 3% normal horse serum (NHS) in 0.2% PBS with Triton X-100 detergent (PBST) at RT. The incubation with the anti-GFP 1.4 nanogold-coupled nanobody was performed overnight at 4°C: anti-GFP in PBS with 0.1% PBST 1:100. On the next day, samples were washed three times for 1 h in PBS at RT and post-fixed for 30 min in 2% glutaraldehyde in PBS at RT. After four washes for 10 min in distilled H_2_O at RT, silver enhancement was performed for 4 min in the dark using the Nanoprobes Silver enhancement Kit (Nanoprobes, USA). After incubation, the solution was quickly removed and washed twice with water for a few seconds. After removal of the enhancement solution, 4x10 min washing steps in distilled H_2_O were performed. Subsequently, samples were fixed for 30 min in 2% OsO_4_ in 0.1 M cacodylate buffer (pH 7.2) and washed 1 h in distilled H_2_O. Samples were further washed over night in distilled H_2_O at 4°C. On the next day, dehydration was performed: 5 min 30%; 5 min 50%; 10 min 70%; 2x10 min 95%; 3x12 min 100%; 1x30 min, 1x1.5 h 50% pure EtOH and 50% epoxy resin (Agar100, Plano, Germany) at RT on a shaker. Samples were then incubated overnight in pure epoxy resin at RT on a shaker. On day four, another incubation step took place in pure epoxy resin for 6 h on a shaker and finally samples were transferred to embedding moulds with fresh epoxy resin for polymerization for 2 days at 70°C.

### Stoichiometric conjugation of an anti-GFP nanobody with 1.4 nm gold particle

Anti-GFP nanobody carrying a single ectopic cysteine at its C-terminus (NanoTag Biotechnologies GmbH, Cat# N0301-1mg) was used for conjugation with 1.4 nm mono- maleimide gold particles (Nanoprobes Inc., Cat# #2020-30NMOL). The ∼30 nmol of anti-GFP nanobody was first reduced using 10 mM tris (2-carboxyethyl)phosphine (TCEP) for 30 minutes on ice. Next, the excess of TCEP was removed using a NAP-5 gravity column and the nanobody immediately mixed with lyophilized 120 nmol of mono-maleimide 1.4 nm gold particles. The mixture was incubated for 4 h on ice with sporadic movement. Finally, the excess of unconjugated gold was removed using an Äkta pure 25 FPLC, equipped with a Superdex 75 Increase 10/300 column.

### Sample mounting for Opto-HPF

After dissection, the samples (*Ai32VC cre^+^* or *Ai32KI cre^+^* (both: (*fl/+ cre^+^ or fl/fl cre^+^*)) were mounted upside down (Fig. 4B,C) due to the specific insertion mechanism of the high-pressure freezing machine (HPM)100. The sapphire disc of 6 mm Ø and 0.12 mm thickness (Leica Microsystems, Wetzlar, Germany) was placed into a sample holder middle plate with a rim of 0.2 mm (Leica Microsystems, Wetzlar, Germany). Thereafter, the first 6 mm Ø and 0.2 mm thick spacer ring (Leica Microsystems, Wetzlar, Germany) was placed, forming a cavity. The freshly dissected organ of Corti was then placed into this cavity that was filled with extracellular solution. The dissection and extracellular solutions had the same composition as the solutions used for patch-clamp recordings (see above). The 0.2 mm side of the 6 mm Ø type A aluminum carrier (Leica Microsystems, Wetzlar, Germany) was placed onto the sample firmly. Next, the second spacer ring of the same dimensions as the first one was placed over the carrier, making a 1.02 mm sample enclosure. Finally, the samples were sandwiched between two transparent cartridges (Leica Microsystems, Wetzlar, Germany). The sample sandwich was then flipped 180° during the insertion process allowing the sample to face towards the light source inside the HPM100.

### Setup for stimulation and freezing relay

The HPM100 (Leica Microsystems, Wetzlar, Germany) is equipped with a trigger box and an optical fiber that reaches the freezing chamber inside the machine (Fig. 4A). The HPM100 allows immediate initiation of the freezing process after loading the sample on the cartridge mount and pressing the *process* button of the HPM100. However, the company configuration does not provide a precise temporal control of the freezing process onset immediately after the light stimulation is over. Therefore, we installed an external control for the blue light-emitting diode (LED) stimulation and subsequent freezing process initiation. This setup had three key functionalities: (i) external control of the *stimulus* onset and duration; (ii) precise control of the time point at which the freezing process initiates by interfacing with the optical trigger box (account for *HPM start* and *HPM delays* from the start); (iii) command relay to the *accelerometer*, the *pneumatic pressure sensor* and the *microphone* to detect the mechanical and acustical processes within the HPM till the end of the freezing process (Fig. 4A, Fig. 5).

To have an external control of the irradiance and duration of the light stimulation used for Opto-HPF (Fig. 4B,C), a *LLS-3* LED blue light (A20955; 473 nm) source (Schott and Moritex) was used for stimulation. *LLS-3* allowed the distinction between manual (by intensity control knob, which is maintained at 0) and automated intensity control (through *RSS-232* input). The latter was used with 80 mV selected as the command voltage (see calibration of irradiance at sample below). A *PCI 6221* interface card (37 pins, National Instrument, NI) and a RS232 interface was used to communicate the external LED control box to the computer. This allowed to control the amplitude and duration of the light pulse via the computer interface (Source code 3). A flexible optical fiber (Leica Microsystems, Wetzlar, Germany) transmitted the blue light from the source to the sample inside the freezing chamber of the HPM.

The *START remote port* of the optical trigger box was connected to the HPM100 via the *J3* cable. This allowed to *START* and *PAUSE* the freezing process externally (either manually or automatically). Light pulse duration could be defined manually or automatically via the computer interface to have different light stimulation durations of the specimen.

### Irradiance measurements for the HPM100

In order to be able to determine the light irradiance that reaches the sample, the inside of the HPM freezing chamber was replicated in an in-house workshop. In this chamber copy, the sample carriers with the upper half-cylinder are included, as well as the LED light source. The exact angle and distances of the original machine are fully replicated based on the technical drawing kindly provided by Leica Microsystems.

The radiant flux (*Φ_e_*) measurement involves two main custom-made components, a mechano-optical arrangement and an optical detector. As optical detector, we used a combination of a bare photodiode (First Sensor, PS100-6 THD) and an operational amplifier circuit (operational amplifier: Burr Brown, OPA 637). It incorporates negative voltage bias over the photodiode and a low noise setting due to relative strong current feedback. The photodiode was covered with a neutral density (ND) filter (5% transmission) and brought as close as possible to the sample plane (Fig. 4D, upper panel). The ND filter was necessary to do not drive the circuit into saturation. The whole detector arrangement was calibrated with a laser light source emitting at 488 nm, which corresponds to the center wavelength of the LED source. In more detail, the calibration was performed with an expanded beam that fits well on the active area of the photodiode and a ND filter with calibrated transmission (active area: 10 x 10 mm). The radiant flux can be calculated from the detector output voltage (*U*) with the linear equation:

*Φ_e_(U)=0.67+0.64 [mW/V] *U [V]*20*

The light distribution measurements were performed by imaging scattered excitation light in the sample plane (Fig. 4C). For imaging, we used two Achromates (*f = 50 mm*) in a configuration with a magnification factor of 1:1 and a CCD camera (IDS, UI-3250ML-M- GL) as detector (Fig. 4D, lower panel). The spatial irradiance distribution (Fig. 4F) is derived by transferring the gray values of the image (Fig. 4E; source code 4) into a radiometric magnitude. Here, we determined the intensity of each pixel (*E_e_*/*p*). First, the intensity per gray value (*gv*) increment was calculated by normalizing the sum of the gray values (- background) in the imaged area to the measured radiant flux (*Φ_e_*). By multiplying the *gv* of each pixel, we determined the intensity per pixel

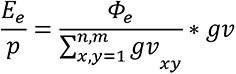

The radiant flux at the sample was measured to be 37.3 mW with a peak irradiance of 6 mW/mm^2^ at the center of the chamber (Fig. 4F).

### Installing additional sensors at the HPM100

The HPM100 initiates the freezing process directly after loading the sample (for further details, refer to https://www.leica-microsystems.com) but does not precisely trigger and monitor the freezing process on an absolute millisecond time scale regarding the externally initiated *HPM start*. The internal pressure and temperature sensors of the HPM100 offer a freezing curve for each sample with the precise values for temperature and pressure development inside the freezing chamber. These values are stored on a USB stick and are available as an excel file. This internal recording only starts when the internal pressure measured in the freezing chamber reaches 65 bar and, therefore, it does not provide the absolute time elapsed between *HPM start* (t = 0) and the time point when the sample is frozen. We incorporated three external sensors, (i) an *accelerometer,* (ii) a *microphone,* and (iii) a *pneumatic pressure sensor* (Fig. 4, Fig. 5, description see below). These additional sensors allowed us to calculate the absolute time scale from the *HPM start* till the sample reaches 0°C (assumed as “frozen”) for each shot (Fig. 5).

### Accelerometer

The accelerometer from Disynet GmbH (Germany) was externally installed under the HPM100 in order to detect vibration caused during the whole process from sample insertion till pressure release.

### Microphone

A microphone (MKE2, Sennheiser electronic GmbH & Co, Germany) was installed inside the machine close to the freezing chamber in order to detect acoustic signal changes during the process from sample insertion till pressure release.

### Pneumatic pressure sensor

The pneumatic pressure sensor (pressure sensor type A-10, WIKA, Germany; different from the HPM100 internal pressure sensor sitting inside the freezing chamber) was installed below the pneumatic needle valve, which opens at 7 bar (pneumatic pressure valve) to regulate the LN_2_ entry in the freezing chamber. The pneumatic pressure sensor detects the pneumatic pressure build-up and reveals the exact time point when the valve opens and the pneumatic pressure drops. This sensor is controlled by the same external control unit that also triggers the start of the HPM and was proven to be the most reliable readout to correlate freezing of the samples to the stimulation.

The absolute time scale (in ms) was correlated to the typical pressure curve inside the freezing chamber. This curve shows a pressure build-up (critical pressure of 1700 bar), followed by a plateau during freezing, and finalized by a rapid pressure drop. The temperature curve, detected by the internal thermal sensor, on the other hand shows a steady drop of the temperature (Fig. 5). These curves obtained from the internal sensors (pressure and temperature inside the freezing chamber) were correlated to the signals obtained from the three external sensors, whereby the pneumatic pressure sensor delivered a clear characteristic signal at the beginning of the pressure build-up (Fig. 5C; source code 5). The recordings from the external sensors start when *HPM start* (t = 0) (Fig. 5).

### Opto-HPF and freezing procedure

When the HPM was ready for freezing (showing “ready for freezing” on the HPM display), the freezing process was halted externally by pressing *PAUSE* on the trigger box (Leica Microsystems, Wetzlar, Germany). Subsequently, the sample sandwich was mounted as described above (Fig. 4B,C) and inserted into the HPM. By pressing the *START* (*HPM start*: t = 0) command (undoing the *PAUSE* command), the process was resumed: the sample was stimulated for the duration chosen with the installed computer interface and the freezing proceeded to completion.

Several factors need to be considered to precisely calculate the time point when the sample is frozen after *HPM start*. This delay, referred as *T_HPM delay from START_*, is a sum of the delays caused by LN_2_ compression and entry, mechanical processes inside the machine (e.g. placing the cartridge in the freezing chamber, valve openings) and specimen freezing. It is the full time it requires from initiating the freezing process by pressing START till the time point, when the sample is frozen. A good quality of freezing requires a steep pressure-increase and rapid temperature-drop inside the freezing chamber. Since these parameters vary for each shot, it is critical to measure them to accurately calculate *T_HPM delay from START_*. Furthermore, the time for pressurizing LN_2_ strongly relies on mechanical processes inside the machine and therefore is variable, too. According to the HPM100 manual, the time to pressurize the LN_2_ (*T_N2 pressurized_*) is around ∼400 ms after *HPM start*. Once LN_2_ reaches the required pressure, the pneumatic needle valve opens to let LN_2_ inside the freezing chamber. The external pneumatic pressure sensor detects these changes in pneumatic pressure outside the freezing chamber (Fig. 5A,B). The recorded pneumatic pressure curve shows a small dip (Fig. 5C, inset, asterisk) before the final steady increase and sudden drop. This small dip in the pressure reflects the opening of the valve. This pressure dip recorded by the external sensor can be correlated to the pressure builtup recorded by the internal sensor in the freezing chamber. This correlation sets the absolute time axis and determines the duration of the mechanical delays prior to freezing (*T_mechanics_*). We also account for the time required for the specimen to reach 0°C (*T_specimen at 0_*). The exact temperature at the specimen cannot be monitored (Watanabe et al., 2013b), as the internal *thermal sensor* only provides the information of the temperature at the freezing chamber. In our HPM100 instrument, the freezing chamber reached 0°C (*T_chamber at 0_*) at 5.41 ± 0.26 ms (SD) on average. This parameter was calculated from the summation of *rise time* and *shift* (*p/T*) from 10 test shots, similarly to a previous report (Watanabe et al., 2013b). *Rise time* corresponds to the time required for the pressure to reach 2100 bar, while *shift* p/T describes the time required for the temperature to drop below 0°C in relation to the pressure rise. Further delays include the time required for the sapphire disc to cool down (*T_sapphire at 0_*) (0.01 ms, as estimated in (Watanabe et al., 2013b)) and for the sample center to reach 0°C (*T_sample center at 0_*) (1.1 ms, as estimated in Watanabe et al., 2013b). Alltogether, the specimen reaches 0°C in approximately 6.52 ms (*T_specimen at 0_ = T_chamber at 0_ + T_sapphire at 0_ + T_sample center at 0_*). This time might be an overestimation since we assume that the sample does not cool during the first ms after the chamber is filled with LN_2_, as stated previously (Watanabe et al., 2013b). Overall, we estimated the delay from *HPM start* as follow:

*T_HPM delay from START_*

*= T_N2 pressurized_+ T_mechanics_ + T_specimen at 0_*

*= T_N2 pressurized_+ T_mechanics_ +* (*T_chamber at 0_ +T_sapphire at 0_ + T_sample center at 0_*) = 400 + **individually determined per shot** + ∼5.41 + 0.01 + 1.1 (± 0.26 ms due to the variability of *T_chamber at 0_*).

Where *T_N2 pressurized_* = 400 ms, and *T_specimen at 0_* = 5.41 (± 0.26) + 0.01 + 1.1 ms.

*T_mechanics_* ranged between 25.4 to 41.6 ms, and was individually determined for each shot. The *T_HPM delay from START_* for our experiments ranged from 431.92 to 448.12 ms. The onset (StimStart) of a 100 ms light stimulus was set after 390 and 425 ms from *HPM start*. We subtracted StimStart from *T_HPM delay from START_* to obtain the actual stimulation duration before freezing for each shot (Fig. 5D).

Stim = *T _HPM delay from START_* – StimStart

ShortStim = *HPM_delay from START_* – 425 ms

LongStim = *HPM_delay from START_* – 390 ms

The light stimulation duration ranged between 17-25 ms for ShortStim and between 48- 76 ms LongStim.

### Sample processing via freeze substitution, ultrathin sectioning and post-staining

Freeze substitution (FS) was performed in an EM AFS2 (Leica Microsystems, Wetzlar, Germany) according to published work (Chapochnikov et al., 2014; Jung et al., 2015a; Siksou et al., 2007; Vogl et al., 2015). Briefly, the samples were incubated for four days in 0.1% tannic acid in acetone at -90°C. Three washing steps with acetone (1 h each) were performed at -90°C. Then 2% osmium tetroxide in acetone was applied to the sample and incubated at -90°C for 7 h. The temperature was raised to -20°C (5°C/h increment) for 14 h in the same solution. The samples were then incubated at -20°C for 17 h in the same solution. The temperature was further raised automatically from -20°C to 4°C for 2.4 h (10°C/h increment). When the temperature reached 4°C, the samples were washed in acetone three times (1 h each) and brought to room temperature by placing them under the fume hood. Finally, the samples were infiltrated in epoxy resin (Agar 100, Plano, Gemany). The next day, the samples were embedded in fresh 100% epoxy resin and polymerized at 70°C for 48 h in flat embedding moulds.

After trimming, 70 nm ultrathin sections were obtained using a UC7 ultramicrotome (Leica Microsystems, Wetzlar, Germany) with a 35° diamond knife (DiAtome, Switzerland). These sections were used to control for freezing quality and find the region of interest, or to perform pre-embedding immunogold labeling. For ET, semithin 250 nm sections were obtained. Post-staining was performed with 4% uranyl acetate in water or uranyl acetate replacement solution (Science Services, EMS) for 40 min and briefly (< 1 min) with

Reynold’s lead citrate in a closed staining compartment in the presence of NaOH to exclude atmospheric CO_2_ and avoid lead precipitates. After that, grids were washed two times on water droplets with previously boiled and cooled distilled water.

### Transmission electron microscopy and electron tomography

To check pre-embedding immunogold labeling and the quality of the tissue, 2D electron micrographs were taken from 70 nm ultrathin sections at 80 kV using a JEM1011 TEM (JEOL, Freising, Germany) equipped with a Gatan Orius 1200A camera (Gatan, Munich, Germany). In the 250 nm sections, we further controlled for artifact-free tomograms, whereby samples with poor tissue integrity and freezing artifacts at the AZ were excluded. These freezing artifacts were identified by the formation of long filamentous artifacts in the cytoplasm and nucleus of the IHCs. Additionally, only tomograms with a continuous AZ, a clear synaptic cleft and round-shaped SVs were analyzed.

*Electron tomography* was performed as described previously (Chakrabarti et al., 2018; Wong et al., 2014). Briefly, 10 nm gold beads (British Bio Cell/Plano, Germany) were applied to both sides of the stained grids. For 3D, tilt series were acquired at 200 kV using a JEM2100 TEM (JEOL, Freising, Germany) mostly from −60° to +60° with a 1° increment at 12,000× using the Serial-EM software package (Mastronarde, 2005) with a Gatan Orius 1200A camera (Gatan, Munich, Germany). Tomograms were generated using the IMOD package etomo (Kremer et al., 1996).

### Model rendering and image analysis

Tomograms were segmented semi-automatically using 3dmod (Kremer et al., 1996) with a pixel size of 1.18 nm. The presynaptic AZ membrane was defined by the area occupied by a clear postsynaptic density (PSD) as well as a regular synaptic cleft. The AZ membrane of a ribbon synapse was then assigned as a closed object and manually segmented every 15 virtual sections for 5 consecutive virtual sections and then interpolated across the Z-stack. The synaptic ribbons and the presynaptic density were also assigned as closed objects and were manually segmented for the first 10, middle 20 and last 10 virtual sections and then interpolated across the Z-stack using the interpolator tool of 3dmod. Interpolation was corrected manually in each virtual section thereafter.

MP-SVs were defined as the vesicles localized in the first row from the AZ-membrane, with a maximum 50 nm membrane-to-membrane distance vertically to the AZ-membrane (Fig. 6-figure supplement 1A, upper panel) and with a maximum lateral distance (vesicle outer edge) of 100 nm to the presynaptic density (Fig. 6-figure supplement 1A, lower panel) (Chakrabarti et al., 2018; Jung et al., 2015a). The distances of the SVs to the presynaptic density or the AZ membrane were measured using the *“Measure”* drawing mode in IMOD’s GUI “*3dmod*”. RA-SVs were defined as the first row of SVs with a maximal distance of 80 nm from the ribbon surface to the vesicle membrane in each tomogram (Fig. 6-figure supplement 1A, upper panel).

All round vesicles were annotated using a spherical scattered object at its maximum projection in the tomogram, marking the outer leaflet of the vesicles. The diameter of the sphere was adjusted for each vesicle. The vesicles radius (r) were determined automatically (Helmprobst et al., 2015) using the program imodinfo option –p of IMOD software package (Kremer et al., 1996). Then the diameter (D) was computed with D = 2r. All outputs were obtained in nm/tomogram.

### Data analysis

Electrophysiological data was analyzed using the IgorPro 6 software package (Wavemetrics; RRID:SCR_000325), Patchers Power Tools (RRID:SCR_001950) and a custom-written script (source code 2). Evoked photocurrents and photodepolarizations were estimated from the peak of current and depolarization, respectively, following the light pulse. Time to peak was calculated from the onset of the light stimulus to the peak of the photodepolarization. EPSCs amplitude and charge was computed from the onset of the light pulse until the end of the release. EPSCs latency was calculated from light- pulse_onset_ until EPSC_onset_ (corresponding to avg. baseline ± 4 SD) and the time of return to baseline was estimated by EPSC_offset_-EPSC_onset_.

Confocal sections were visualized using the FIJI software (Schindelin et al., 2012; RRID:SCR_002285). The analysis of ribbon number using the *z*-projections of the stacks was performed in the IMARIS software using custom plug-ins (Source code 1) of IMARIS (RRID:SCR_007370), whereby the number of ribbons within a region of interest (ROI) was obtained using the Spots function. The average ribbons per IHCs was calculated by dividing the number of spots detected by the number of IHCs for each ROI.

For every dataset, the number of replicates (n) and number of animals (N) are indicated in the figure legend. Data sets were tested for normal distribution (Saphiro-Wilk test) and equality of variances (Brown-Forsythe test). For data sets following a normal distribution and with equality of variances, we used parametric statistical tests (one-way ANOVA and two-way ANOVA), followed by a post-hoc test for multiple comparisons (Tukey’s test). For non-parametric data sets, we performed Kruskal-Wallis (KW) tests, followed by Dunn’s test. For the SV diameter quantification (Fig. 7C), we also categorized SV diameters into bins similar to previous studies (Chakrabarti et al., 2018; Hintze et al., 2021). All statistical analyses and graphs were done using IGOR Pro software 6, GraphPad Prism (RRID:SCR_002798) version 9 and/or R software (version 4.0.3).

Sample sizes were decided according to typical samples sizes in the respective fields (e.g. electrophysiology, electron tomography). The sample size for each experiment is reported in the main text, figures and figure captions.

### Materials availability statement

All research materials and biological reagents used in this paper are reported in the Materials and Method section. The custom routines and scripts used in the manuscript are provided as Source Codes:

Source Code 1: IMARIS custom plug-ins for the analysis of Figure 1D

**Figure 1.**
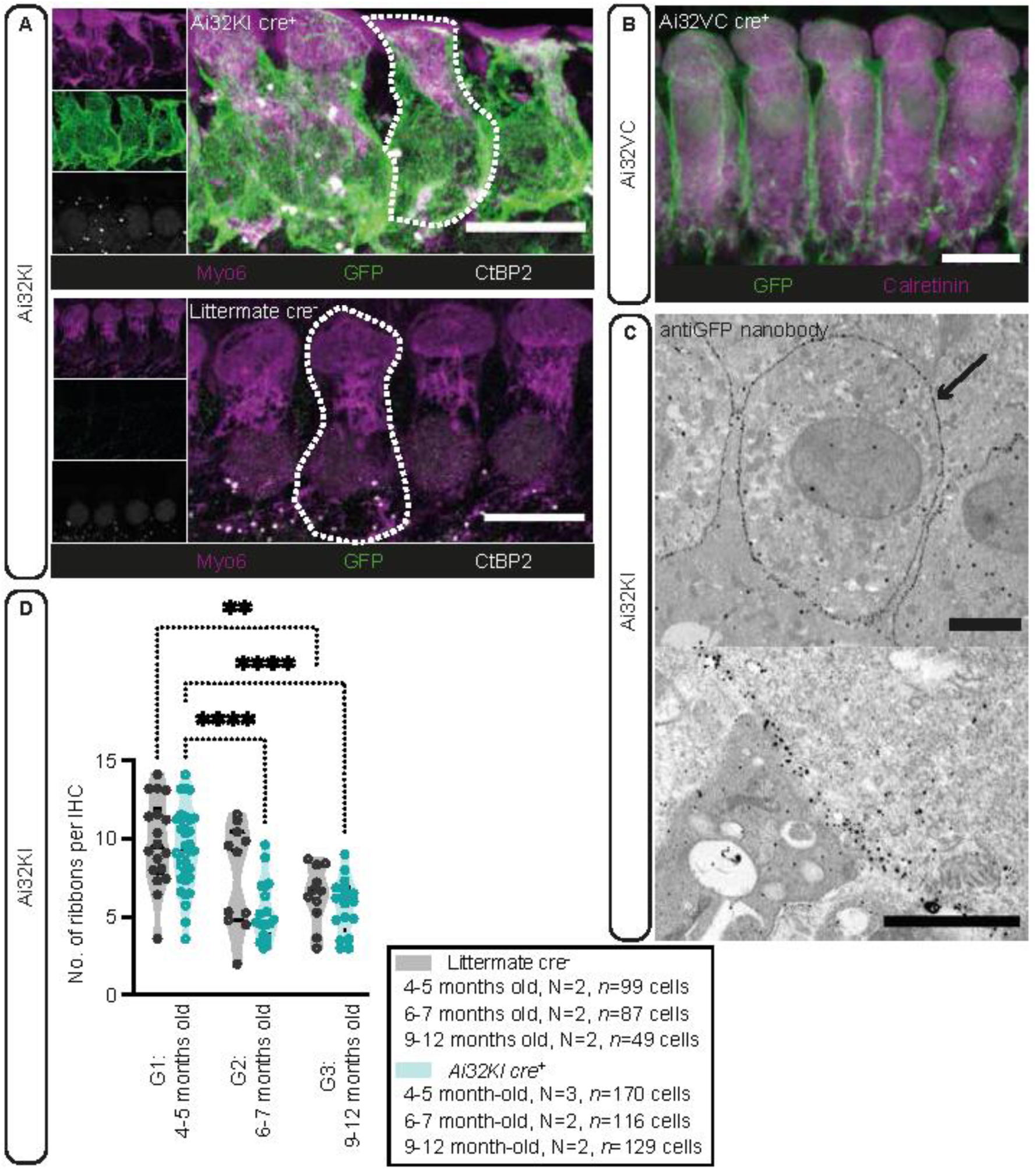
Plasma membrane expression of ChR2 in IHCs. (**A**) IHCs express ChR2 in the plasma membrane. Maximal projection of confocal z-stacks from the apical turn of the anti-GFP labeled organ of Corti from a 4-5 months-old *Ai32KI cre^+^* mice (upper panel) and its littermate control (WT) (lower panel). Myo6 (magenta) was used as counterstaining and CtBP2 labeling shows ribbons at the basolateral zone of IHCs (examples outlined). ChR2 expression (green) is observed along the surface of *Ai32KI cre^+^* IHCs (upper panel) but not in WT IHCs (lower panel). Scale bar, 10 μm. **(B)** Maximal intensity projection of *Ai32VC cre^+^*IHCs expressing ChR2-EYFP construct. Calretinin immunostaining was used to delineate the IHC cytoplasm. Scale bar, 10 µm. (**C**) Pre-embedding immunogold labeling for electron microscopy performed with a gold-coupled anti-GFP nanobody recognizing the EYFP of the ChR2 construct. A clear localization at the plasma membrane of IHCs is visible (arrow, upper panel). Scale bar, 2 μm. Magnification of the membrane labeling is shown in the lower panel. Scale bar, 1 μm. **(D)** The average number of ribbons per IHC is similar between ChR2-expressing IHCs and WT IHCs. Both *Ai32KI cre^+^* and WT mice showed a comparable decrease in the number of ribbons with age (* p < 0.05; ** p < 0.01; **** p < 0.0001; two-way ANOVA followed by Tukey’s test for multiple comparisons). Violin plots show median and quartiles with data points overlaid.

Source code 2: Igor Pro custom-written analysis (*OptoEPSCs*) of light-evoked EPSCs related to Figure 3C-F.

**Figure 2.**
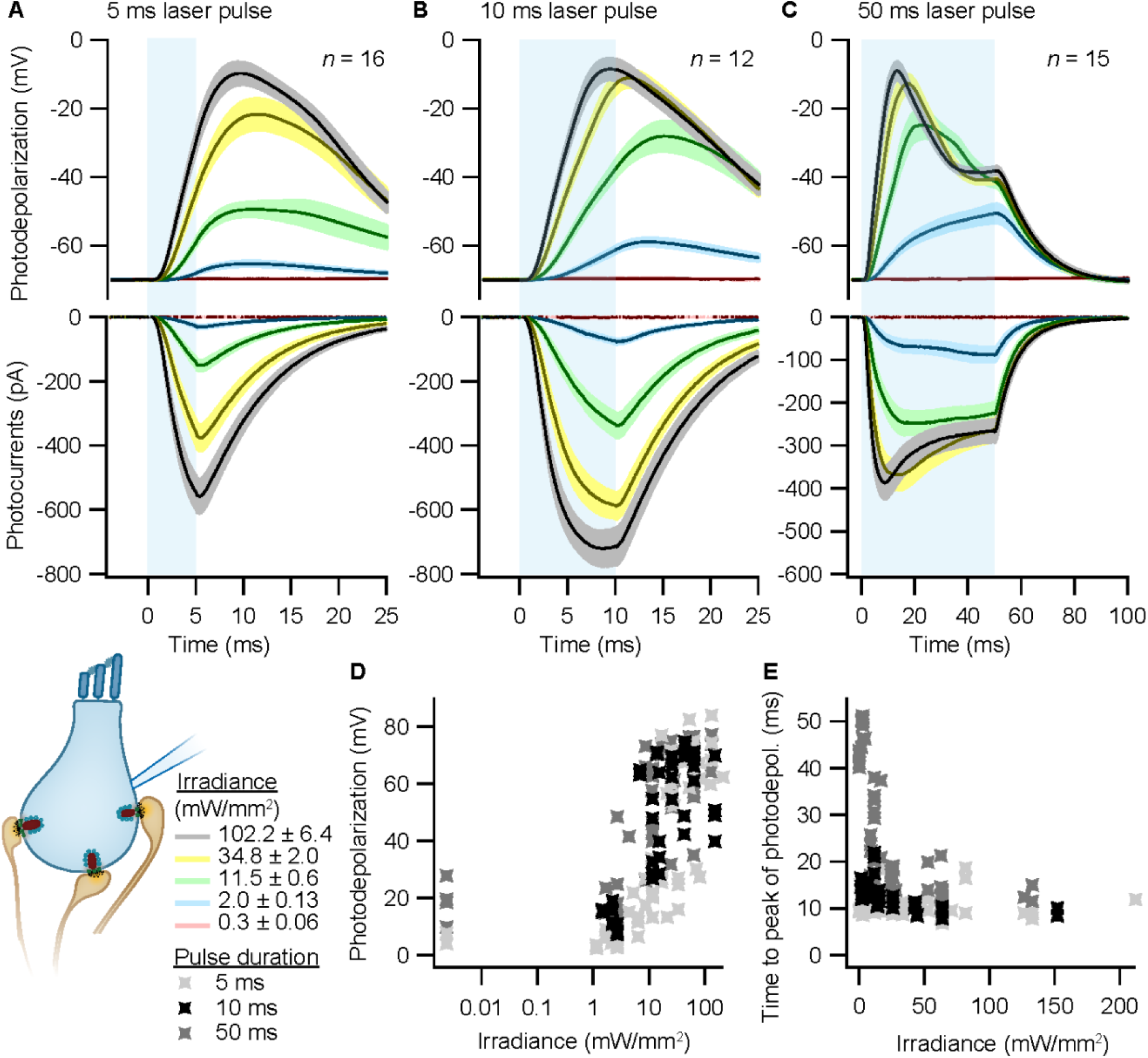
Optogenetic depolarization of IHCs. IHCs expressing ChR2 (*Ai32VC cre^+^* and *Ai32KI cre^+^*) were optogenetically stimulated by 473 nm light pulses of increasing irradiance (mW/mm^2^). **(A-C)** Average photocurrents (lower panel) and photodepolarizations (upper panel) of patch-clamped IHCs during 5 ms (A, *n* = 16, *N_animals_* = 7), 10 ms (B, *n* = 12, *N_animals_*= 4) and 50 ms (C, *n* = 15; *N_animals_* = 6) light pulses of increasing irradiances (color coded). Mean is displayed by the continuous line and ± SEM by the shaded area. **(D-E)** Peak of photodepolarization (D) and time to peak (E) obtained for increasing irradiances of different lengths (light gray 5 ms, black 10 ms, dark gray 50 ms).

**Figure 3.**
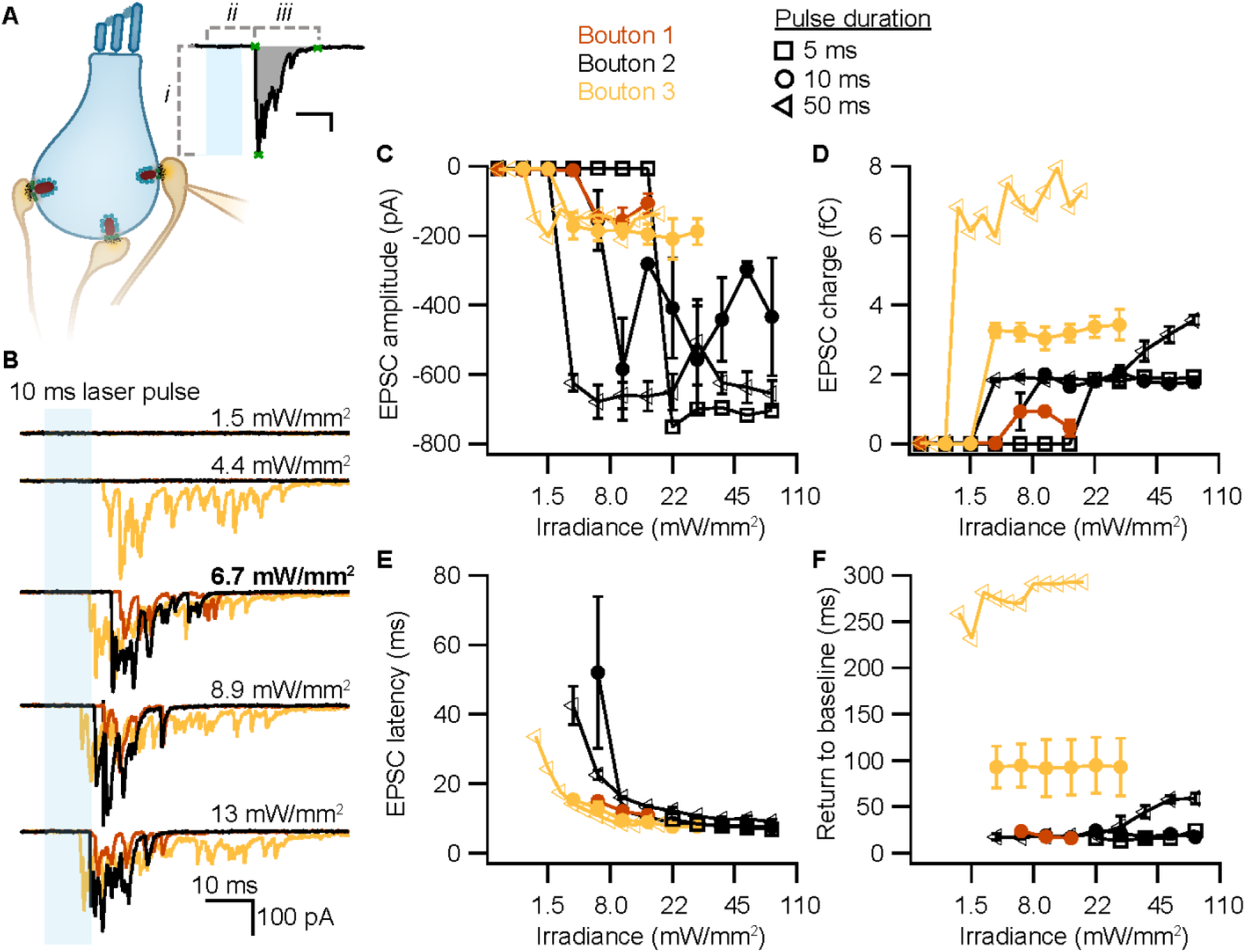
Triggered exocytosis at individual ribbon synapses. **(A)** Excitatory postsynaptic currents (EPSCs) upon the optogenetic stimulation of *Ai32VC cre^+^*IHCs were recorded using whole-cell patch clamp of the contacting bouton. The response was quantified in terms of amplitude (*i*), charge (gray area), latency (*ii*) and return to baseline (*iii*) (Source code 2). Scale bar as in panel B. **(B)** Recorded EPSCs from three different postsynaptic boutons (different colors) in response to increasing light intensities. **(C-F)** Amplitude (C), charge (D), latency (E) and return to baseline (F) of the light triggered EPSCs to different pulse durations (5 ms squares; 10 ms circles; 50 ms triangles).

Source code 3: MATLAB scripts (*HPMacquire*) for the computer interface to control the light pulse for Opto-HPF. Related to Figure 4A

**Figure 4.**
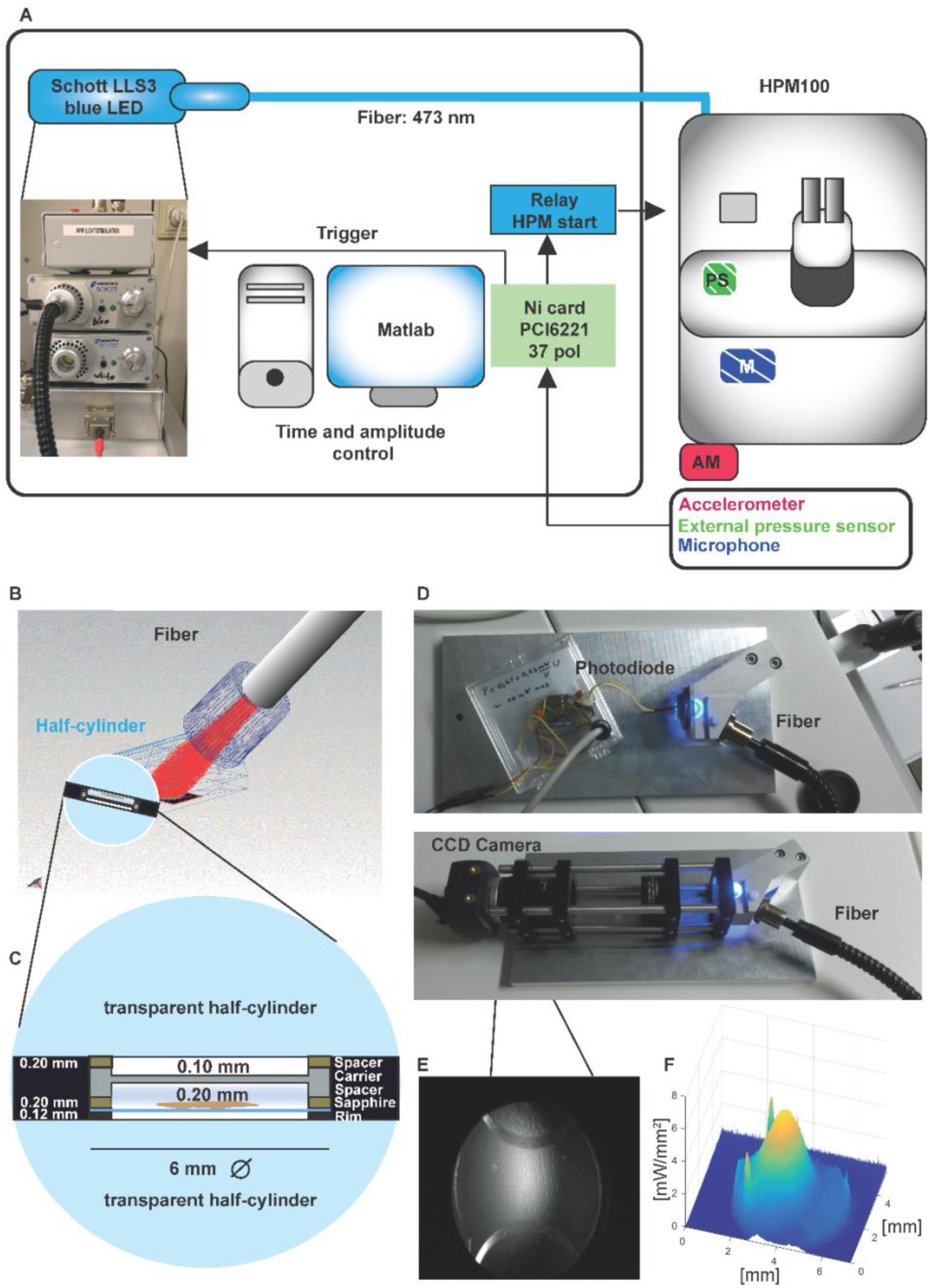
Opto-HPM setup and irradiance calculation in the HPM. **(A)** Simplified illustration depicting the components of the external setup installed to control the light stimulation (irradiance and duration), to determine the time point of the freezing onset and command relay (blue, control unit) to the sensors (accelerometer (red, outside, AM), microphone (patterned blue, inside, M), pneumatic pressure sensor, green, inside, PS) to initiate the mechanical sensing process of the HPM100 (Source code 3). **(B)** Fiber – cartridge arrangement in the HPM100 with the fiber at an angle of 60° to the upper half-cylinder: Sample plane: black, fiber: gray, Light rays: red. Mechanical components of HPM100 are not shown. **(C)** Sample loading scheme **(D)** Rebuild chamber to enable irradiance calculation**. (E)** CCD image of the photodiode in the sample plane. **(F)** The spatial irradiance distribution with a peak irradiance of ∼6 mW/mm^2^ at 80% intensity of the LED was calculated using a self-written MATLAB routine *intensityprofilcalculator.m* (Source code 4). Depicted are pixel values in irradiance.

Source code 4: MATLAB script (*Intensityprofilecalculator*) for the analysis of the irradiance in Figure 4E.

Source code 5: MATLAB scripts (*HPManalyse*) for the alignment of the data obtained from the Opto-HPF sensors. Related to Figure 5C.

**Figure 5.**
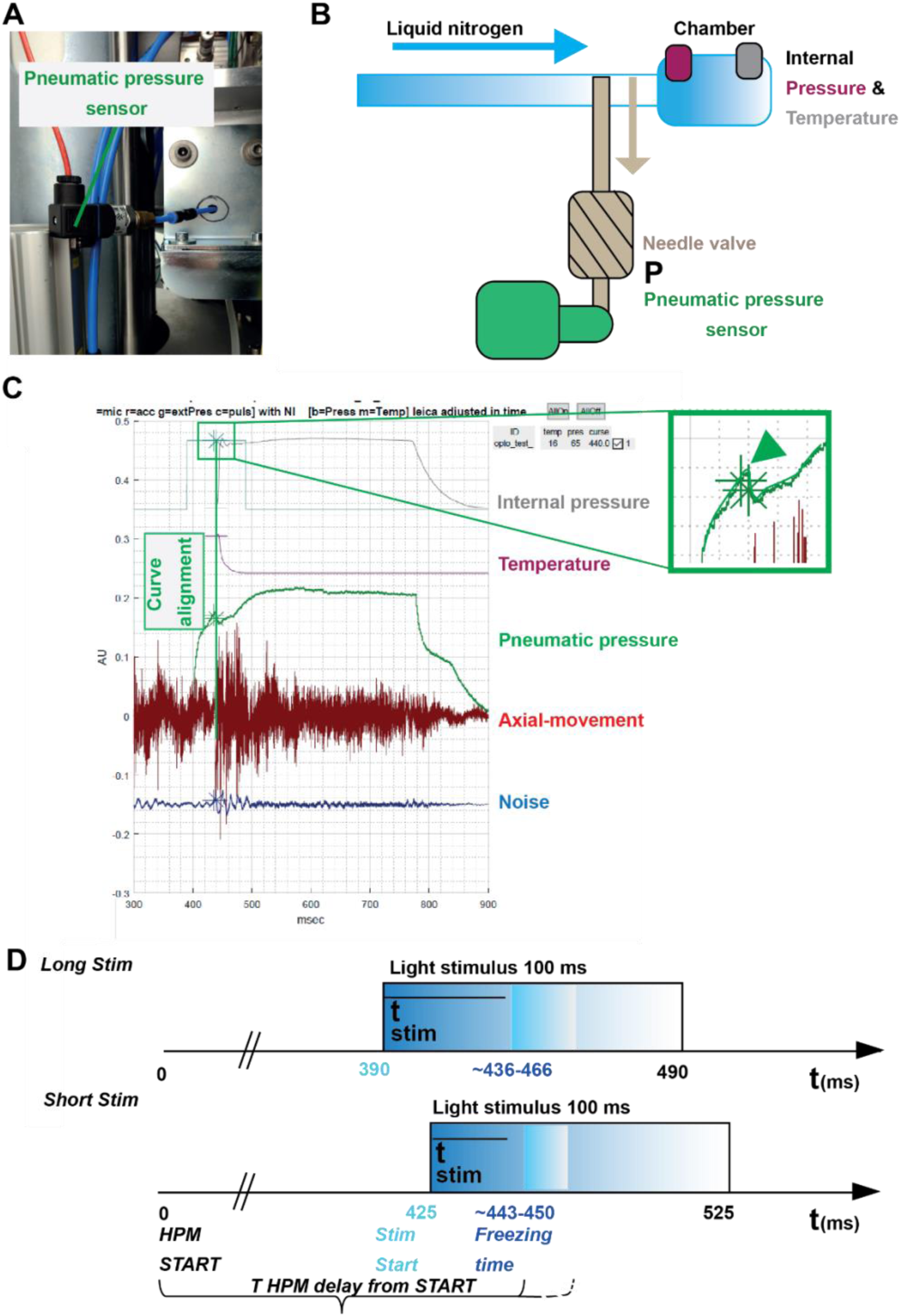
Correlating the sensor signals to internal pressure and temperature measured inside the HPM. **(A)** Pneumatic pressure sensor inside the HPM100. **(B)** Scheme of the localization of the pneumatic pressure sensor below the pneumatic needle valve, which allows LN_2_ influx in the chamber for freezing. **(C)** Depicted are the curves from the different sensors aligned by using the MATLAB GUI (Source code 5). The curves of the pressure-build up start and temperature corresponding decline are aligned to the steep drop in the pneumatic pressure curve (green arrowhead, inset). **(D)** The outline of the optical stimulation incorporated with HPF: 100 ms light pulse set to a 390 ms delay from HPM start resulted in a stimulation duration of 48-76 ms before freezing (“LongStim”) and that set to a 425 ms delay from HPM start in a stimulation of 17-25 ms (“ShortStim”).

The data files including the numerical data associated with the figures will be made available in osf upon acceptance.

## Results

### Verification of ChR2 expression in inner hair cells and long-term expression of ChR2

First, we verified the expression of ChR2 in IHCs of both mouse lines, *Ai32VC cre^+^* and *Ai32KI cre^+^* (*fl/+ cre^+^* or *fl/fl cre^+^*), using immunofluorescence microscopy (Fig. 1A-C). Alexa-coupled anti-GFP antibodies detected the EYFP-tag of the ChR2 construct and showed a clear expression of the construct at the membrane. For better resolution, we also preformed immunogold electron microscopy (Fig. 1D) with pre-embedding gold- coupled anti-GFP nanobodies. Membrane expression of ChR2 was evenly distributed (Fig. 1D, arrow), confirming that ChR2 was efficiently expressed at the plasma membrane of IHCs without any apparent intracellular accumulation. Overall, these results confirmed that the *Vglut3* promoter efficiently controlled cre-recombination in ∼99% of the analyzed IHCs for both mouse lines similarly to previous studies using *Vglut3 cre* mice (Jung et al., 2015b; Vogl et al., 2016). Therefore, we decided to pool the results from both genotypes in the following sections, but also analyzed and presented the data for each genotype in the Expanded View Figures.

Having confirmed proper ChR2 expression in both lines, in a next step, we analyzed potential long-term effects of ChR2 expression on synaptic organization of IHCs in the Ai32KI line. Using confocal microscopy, we compared ribbon synapse numbers of *Ai32KI cre^+^* IHCs with WT littermate controls at three different age intervals: 4-5 months (G1), 6- 7 months (G2) and 9-12 months (G3) (Fig. 1, Fig. 1-figure supplement 1, all values can be found in Table 2). WT IHCs showed the characteristic decline in the number of ribbon synapses associated with aging (WT G1 = 9.82 ± 0.87 vs WT G3 = 6.39 ± 0.56; p = 0.0077) (Parthasarathy and Kujawa, 2018; Sergeyenko et al., 2013). Comparably, ChR2- expressing IHCs showed a significant decline in the number of ribbon synapses at 6-7 months and 9-12months in comparison to 4-5 months, (*Ai32KI cre^+^* G1 = 8.97 ± 0.48 vs *Ai32KI cre^+^* G2 = 5.324 ± 0.43; p < 0.0001; vs *Ai32KI cre^+^* G3 = 5.69 ± 0.42; p = 0.0005). Importantly, there were no differences in the number of ribbons between ChR2- expressing IHCs and WT from the same age groups. We therefore conclude that ChR2 expression does not alter the number of ribbon synapses arguing against adverse effects such as through a potential chronic ChR2-mediated depolarization.

**Table 2.** Ribbon counts in three different age groups of ChR2-expressing IHCs and their WT controls. Data are presented as mean ± SEM. p-values are calculated by two-way ANOVA followed by Tukey’s test for multiple comparisons. Results of the comparisons between age groups of the same genotype and between genotypes of the same age group are reported. Significant results are indicated with * p< 0.05; ** p<0.05; and **** p< 0.0001.

| | N<br>*animals* | N<br>*ROIs* | *n*<br>*cells* | Ribbon<br>count (*mean*<br>± *SEM*) | Age comparison | | Genotype comparison | |
| --- | --- | --- | --- | --- | --- | --- | --- | --- |
|  |  |  |  |  | p-value | Test | p-value | Test |
| WT G1 | 2 | 12 | 99 | 9.82 ± 0.87 | **, WT G1 vs. WT G3 | | ns, WT G1 vs. *Ai32KI cre*<sup>+</sup> G1 | |
|  |  |  |  |  | 0.0077 | Two-way ANOVA | 0.9124 | two-way ANOVA |
| WT G2 | 2 | 11 | 87 | 7.60 ± 0.99 | ns, WT G2 vs. WT G1 | | ns, WT G2 vs. *Ai32KI cre*<sup>+</sup> G2 | |
|  |  |  |  |  | 0.2045 | two-way ANOVA | 0.1046 | two-way ANOVA |
| WT G3 | 2 | 11 | 65 | 6.39 ± 0.56 | ns, WT G3 vs. WT G2 | | ns, WT G3 vs. *Ai32KI cre*<sup>+</sup> G3 | |
|  |  |  |  |  | 0.8179 | two-way ANOVA | 0.9689 | two-way ANOVA |
| *Ai32KI cre*<sup>+</sup> G1 | 3 | 21 | 170 | 8.97 ± 0.48 | ****, *Ai32KI cre*<sup>+</sup> G1 vs. *Ai32KI cre*<sup>+</sup> G2 | | | |
|  |  |  |  |  | <0.0001 | two-way ANOVA |  |  |
| *Ai32KI cre*<sup>+</sup> G2 | 2 | 19 | 116 | 5.32 ± 0.43 | ns, *Ai32KI cre*<sup>+</sup> G2 vs. *Ai32KI cre*<sup>+</sup> G3 | | | |
|  |  |  |  |  | 0.9968 | two-way ANOVA |  |  |
| *Ai32KI cre*<sup>+</sup> G3 | 2 | 17 | 129 | 5.69 ± 0.42 | ****, *Ai32KI cre*<sup>+</sup> G3 vs. *Ai32KI cre*<sup>+</sup> G1 | | | |
|  |  |  |  |  | 0.0005 | two-way ANOVA |  |  |

### Depolarization of IHCs using optogenetics

To validate optogenetic stimulation of IHCs, we performed perforated patch-clamp recordings of ChR2-expressing IHCs and applied 473 nm light pulses of different durations and intensities. Evoked photocurrents and photopotentials were measured in voltage-clamp and current-clamp mode, respectively. We employed TEA-Cl and Cs^+^ in the bath solution in order to partially block K^+^ channels and facilitate photodepolarization. Compared to other compositions (Jaime Tobón, 2015), we found 20 mM TEA-Cl and 1 mM Cs^+^ to support both sufficient amplitudes and acceptable decay kinetics of photodepolarization. Under these conditions, strong short light pulses of 5 ms depolarized the cell by more than 50 mV (i.e. going from a holding potential of -84 mV up to -30 mV; Fig. 2A). Longer light pulses of 10 and 50 ms caused stronger depolarizations even at low irradiances (Fig. 2B,C,D), even though the peak of photodepolarization was reached with considerable delays (Fig. 2E). On average, the peak was reached within 10-20 ms after the onset of the light pulse for irradiances above 1 mW/mm^2^ (Fig. 2E). Higher irradiances decreased the time to peak of the photodepolarization for all stimulation durations. Light pulses of 10 ms at 7-16 mW/mm^2^, corresponding to the irradiance recorded at the HPM machine (Fig. 4E-F), could depolarize the IHC by 27-65 mV (Fig. 2D). Notably, IHCs were photodepolarized by 20 mV within the first 3 to 10 ms of the light pulse, which presumably suffices to trigger vesicle release on IHCs (assuming a resting potential of -58 mV and based on release thresholds reported (Goutman and Glowatzki, 2007; Özçete and Moser, 2020). The results obtained from both lines were comparable, as shown in Figure 2-figure supplement 1.

To test whether our stimulation paradigms would prompt neurotransmitter release, we performed whole-cell patch-clamp recordings from individual afferent boutons contacting ChR2-expressing IHCs. Light pulses of low intensities and short duration were sufficient to trigger release from individual AZs, as proven by excitatory postsynaptic currents (EPSCs) recorded from three boutons contacting different IHCs (Fig. 3A). Consistent with previous experiments employing K^+^ or voltage-clamp stimulation of IHCs (e.g. Chapochnikov et al., 2014; Glowatzki and Fuchs, 2002; Goutman and Glowatzki, 2007), we found variable amplitudes of individual EPSCs. The maximum amplitude of the evoked EPSCs varied between different boutons (from -200 pA to -700 pA), but remained fairly similar for one individual bouton regardless of the light pulse duration (Fig. 3B). In contrast, light pulse duration had a major impact on the duration of the evoked EPSCs (i.e. the duration of exocytosis; Fig. 3C) and consequently, on the total charge transfer (Fig. 3D). In response to a 50 ms stimulation, the evoked release lasted three times longer than in response to a 10 ms light pulse and could reach up to double the initial charge transfer. In line with the recorded IHC photodepolarization, longer light pulses required lower intensities to trigger a response; 4 to 6 mW/mm^2^ were sufficient for 50 ms light pulses. Moreover, as expected from the photodepolarization, EPSCs latency decreased with increasing irradiances (Fig. 3E). Finally, we quantified the maximal EPSCs charge transfer at 20 ms (Q_20ms_) and 50 ms (Q_50ms_) after the onset of the light pulses. These time points reflect phasic RRP release (20 ms) and sustained release (50 ms) of IHC ribbon synapses (Johnson et al., 2017; Michalski et al., 2017; Moser and Beutner, 2000). For light stimulations of 6-7 mW/mm^2^, Q_20ms_ ranged from 236 pC up to 1300 pC while Q_50ms_ ranged from 850 to 2450 pC. Importantly, the first recording of all three boutons showed substantial release exceeding 400 pC at these time points. The return to baseline differed among the 3 boutons and for pulse duration (Fig. 3F), but neurotransmitter release lasted for at least 15 ms. These electrophysiological findings demonstrate that the chosen stimulation paradigms are sufficient to trigger phasic and sustained SV exocytosis in ChR2-expressing IHCs.

### Developing a method to combine optogenetic stimulation with precisely timed freezing

To correlate structure and function, we performed Opto-HPF, followed by freeze-substitution (FS) and subsequent ET (Fig. 4-figure supplement 1). The commercial HPM100 comes with limitations: the light stimulation duration cannot be set precisely and the precise time of freezing is not provided. Therefore, groups had previously already modified the HPM100 with custom-made settings adapted to the needs of central synapses (Watanabe et al., 2013a). Applying this method to a sensory synapse, which does not operate with all-or-nothing action potential stimulation, we established a more general framework for Opto-HPF using the HPM100.

### Setup for stimulation and freezing relay

First, we determined the irradiance that reaches the sample with a re-built chamber-copy equipped with an optical fiber. The radiant flux at the sample was measured to be 37.3 mW with a peak irradiance of 6 mW/mm^2^ where the samples are positioned (Fig. 4F). This irradiance is in accordance with the irradiance values that led to a sufficient depolarization of IHCs to trigger exocytosis in our cell-physiological experiments. With our custom-made setup, we controlled the irradiance, stimulus duration and the coupling of stimulus onset with the freezing of the specimen (Source code 3).

Our three incorporated external sensors, (i) an *accelerometer,* (ii) a *microphone,* and (iii) a *pneumatic pressure sensor* (Fig. 4, Fig. 5) allowed us to calculate for each shot the absolute time scale from the *HPM start* till the specimen is reaching 0°C (*T _HPM delay from START_*, Fig. 5).

The pneumatic pressure sensor was located directly at the pneumatically steered needle valve in front of the freezing chamber (Fig. 5A-C). In contrast to the other sensors, the pneumatic pressure sensor provided a reliably signal of the moment when the needle valve opened to allow influx of pressurized LN_2_ into the freezing chamber (Fig. 5C; green curve; the green arrowhead points out the time point when the needle valve opens). We used this point to align the curves obtained from the internal sensors, which show the internal pressure buildup (Fig. 5C, gray curve) and the gradual temperature decline (Fig. 5C, purple curve) inside the freezing chamber after the opening of the needle valve. We set the onset (StimStart) of a 100 ms light stimulation between *HPM start* (t = 0) and *T _HPM delay from START_* (calculated for each shot; Fig. 5D).

To obtained short stimulations (ShortStim), StimStart was set at 425 ms. Based on the correlation of the pneumatic pressure sensor curve with the internal pressure and temperature curves, the samples were stimulated during ∼17 to ∼25 ms before the freezing onset (Fig. 5D, lower panel). To obtain longer stimulations (LongStim), StimStart was set at 390 ms, which resulted in light stimulation durations from ∼48 to ∼76 ms before the freezing onset (Fig. 5D, upper panel).

### First ultrastructural analysis of optogenetic stimulated IHC ribbon synapses, coupling structure to function

We analyzed the ultrastructural changes upon precise optical stimulation of ChR2- expressing IHCs (*Ai32VC cre^+^* and *Ai32KI cre^+^*). Our two stimulation paradigms (ShortStim and LongStim, Fig. 6C and D) aimed to capture two functional states of exocytosis at IHCs. A ShortStim (∼17-25 ms) might reflect the changes after/during RRP release, while a LongStim (∼48-76 ms) might reflect sustained exocytosis (Moser and Beutner, 2000; Rutherford and Roberts, 2006). We included two controls (i) B6J under light stimulation (B6J Light; Fig. 6A) and (ii) ChR2-expressing IHCs (*Ai32VC cre^+^* and *Ai32KI cre^+^*) without any light stimulation (ChR2 NoLight; Fig. 6B). For the light control on B6J, we chose a LongStim protocol assuming that potential direct light effects are strongest with the longer exposure. Table 3 includes the number of ribbons and animals included from each genotype in the analysis.

**Figure 6.**
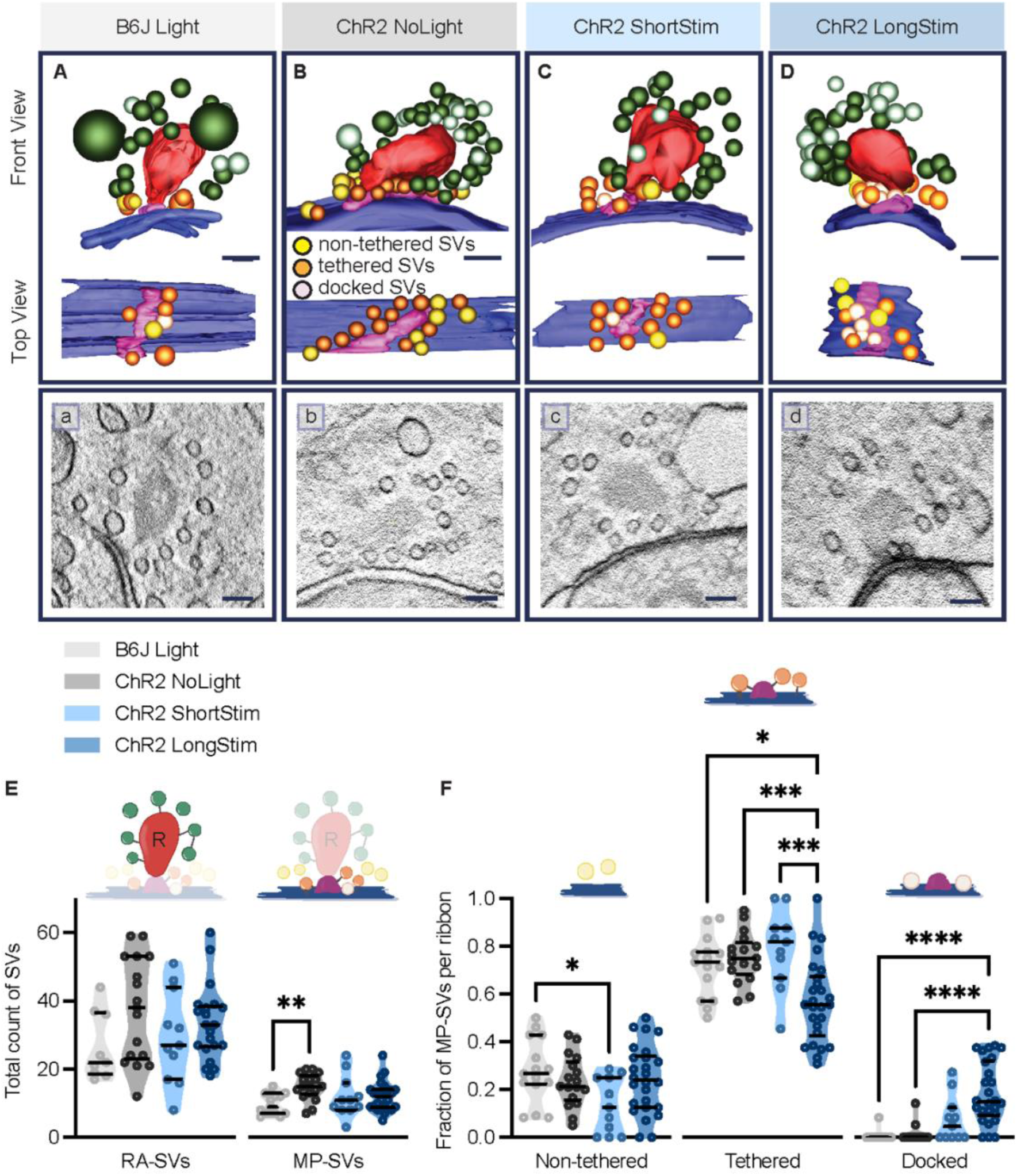
Functional AZ states differ in their morphologically defined vesicle pools. Representative tomographic 3D reconstructions of **(A)** B6J Light**, (B)** ChR2 NoLight**, (C)** ChR2 ShortStim and **(D)** ChR2 LongStim displayed in both front view (upper part of the panel) and top view (lower part of the panel). **(a-d)** Corresponding virtual sections of A-B. The AZ membrane is shown in blue, presynaptic density in pink, ribbons in red, MP-SVs (non-tethered in yellow, tethered in orange and docked in light pink), RA-SVs (green, light green). Magnification 12,000x; scale bars, 100 nm. **(E)** Total count of SVs per pool (RA- and MP-SV pools), per ribbon. **(F)** The fraction of non-tethered, tethered and docked MP-SVs per ribbon. Data are presented in mean ± SEM. *p < 0.05, **p < 0.01, ***p < 0.001 and ****p < 0.0001. Statistical test: one-way ANOVA followed by Tukey’s test (parametric data) and KW test followed by Dunn’s test (non-parametric data). MP-SV pool: B6J Light: *n* = 15 ribbons, *N_animals_* = 2; ChR2 NoLight: *n =* 17 ribbons *N_animals_* = 4; ChR2 ShortStim: *n* = 11 ribbons, *N_animals_* = 1; ChR2 LongStim: *n =* 26 ribbons, *N_animals_*= 4. RA-SV pool: B6J Light: *n* = 9 ribbons, *N_animals_* = 1; ChR2 NoLight: *n =* 17 ribbons *N_animals_* = 4; ChR2 ShortStim: *n* = 11 ribbons, *N_animals_* = 1; ChR2 LongStim: *n =* 21 ribbons, *N_animals_* = 3.

**Table 3.** List of SV parameters showing the mean ± SEM values, N, n, *p*-values and the statistical tests applied. Data are presented as mean ± SEM. Data was tested for significant differences by one-way ANOVA followed by Tukey’s test (parametric data) or KW test followed by Dunn’s test (non- parametric data). Significant results are indicated with * p< 0.05; ** p<0.05; and **** p< 0.0001.

|  | B6J<br>Light | ChR2<br>NoLight | ChR2<br>ShortStim | ChR2<br>LongStim | Adjusted p-value | Test |
| --- | --- | --- | --- | --- | --- | --- |
| *N*<sub>animals</sub> | 2 | 4 | 1 | 4 |  |  |
| *n*<sub>ribbons</sub> Total | 15 | 17 | 11 | 26 |  |  |
| *n*<sub>ribbons</sub><br>Ai32VC |  | 9 | 11 | 15 |  |  |
| *n*<sub>ribbons</sub><br>Ai32KI |  | 8 | 0 | 11 |  |  |
| MP-SVs<br>count | 9.73<br>± 0.796 | 14.88<br>± 0.935 | 12.00<br>± 1.844 | 11.96<br>± 0.833 | B6J Light vs. ChR2 NoLight<br>0.0028 | KW Test -<br>Dunn's test |
| Fraction of<br>non-tethered<br>SVs | 0.28<br>± 0.035 | 0.23<br>± 0.026 | 0.14<br>± 0.035 | 0.25<br>± 0.028 | B6J Light vs. ChR2 ShortStim<br>0.0313 | ANOVA -<br>Tukey's test |
| Fraction of<br>tethered SVs | 0.72<br>± 0.033 | 0.75<br>± 0.025 | 0.79<br>± 0.049 | 0.57<br>± 0.034 | B6J Light vs. ChR2 LongStim<br>0.0173<br>ChR2 LongStim vs. ChR2 NoLight<br>0.0009<br>ChR2 LongStim vs. ChR2<br>ShortStim<br>0.0006 | ANOVA -<br>Tukey's test |
| Fraction of<br>docked SVs | 0.01<br>± 0.006 | 0.01<br>± 0.009 | 0.08<br>± 0.029 | 0.18<br>± 0.025 | B6J Light vs. ChR2 LongStim<br><0.0001<br>ChR2 LongStim vs. ChR2 NoLight<br><0.0001 | KW Test -<br>Dunn's test |
| Distance of<br>SV to the<br>membrane | 22.78<br>± 1.08 | 23.67<br>± 0.776 | 19.41<br>± 1.263 | 19.61<br>± 0.876 | ChR2 LongStim vs. ChR2 NoLight<br>0.0007<br>ChR2 NoLight vs. ChR2<br>ShortStim<br>0.0105 | KW Test -<br>Dunn's test |
| Distance of<br>SV to the PD | 31.88<br>± 2.144 | 37.61<br>± 1.622 | 32.96<br>± 2.192 | 30.13<br>± 1.509 | ChR2 LongStim vs. ChR2 NoLight<br>0.0001 | KW Test -<br>Dunn's test |
| Diameter of<br>SVs | 49.00<br>± 0.442 | 48.67<br>± 0.25 | 49.17<br>± 0.352 | 48.05<br>± 0.303 |  | KW Test -<br>Dunn's test |
| *N*<sub>animals</sub> | 1 | 4 | 1 | 3 |  |  |
| *n*<sub>ribbons</sub> Total | 9 | 17 | 11 | 21 |  |  |
| *n*<sub>ribbons</sub><br>Ai32VC |  | 9 | 11 | 10 |  |  |
| *n*<sub>ribbons</sub><br>Ai32KI |  | 8 | 0 | 11 |  |  |
| RA-SVs<br>count | 26.44<br>± 3.296 | 37.35<br>± 3.683 | 29.45<br>± 4.059 | 33.33<br>± 2.372 | n.s | ANOVA -<br>Tukey's test |

We started our ultrastructural analysis with the determination of the total count of MP-SVs and RA-SVs (Fig. 6-figure supplement 1A). We found no alterations in the size of the MP- SV and RA-SV pools among the various conditions and controls (Fig. 6E), except for a larger MP-SV pool in ChR2 NoLight compared to B6J Light. This might reflect different proportions of ribbon-type AZs contained in the tomograms obtained from 250 nm sections that do not always include the full AZs.

### Enhanced SV docking in correlation to stimulus duration

Next, we performed in-depth analysis of MP-SV sub-pools among the various conditions (Fig. 6F). As the full inclusion of the ribbon is relatively rare in 250-nm sections (Fig. 6A- D), we compared the fractions of non-tethered, tethered and docked MP-SVs as done previously (Chakrabarti et al., 2018). The fraction of non-tethered SVs decreased after a short light pulse (B6J Light = 0.28 ± 0.03, ChR2 NoLight = 0.23 ± 0.03, ChR2 ShortStim = 0.13 ± 0.03), while the fraction of tethered SVs decreased upon a long stimulation (B6J Light = 0.71 ± 0.03, ChR2 NoLight = 0.75 ± 0.02, ChR2 ShortStim = 0.79 ± 0.05, ChR2 LongStim = 0.57 ± 0.03). Values and information about statistics can be found in Table 3 and a separate quantification of the used ChR2 mice can be found in Fig. 6-figure supplement 1.

The fraction of morphologically docked SVs increased upon optogenetic stimulation (Fig. 6F), being more prominent upon a long stimulation (B6J Light = 0.005 ± 0.005, ChR2 NoLight = 0.01 ± 0.008, ChR2 ShortStim = 0.076 ± 0.03, ChR2 LongStim = 0.18 ± 0.02). We conclude that optogenetic stimulation changes the sub-pools of MP-SVs, with a prominent increase of docked SVs proportional to the stimulation duration and fewer tethered and non-tethered SVs.

### SV distances to the PD and AZ membrane decrease upon long stimulation

It was previously proposed that SVs are recruited to the AZ membrane via tethers, a process that takes place rather close to the presynaptic density (PD) (Chakrabarti et al., 2018; Frank et al., 2010; Vogl et al., 2015). Therefore, we evaluated the distances of all MP-SVs to the AZ membrane and to the PD. We found that the distances of MP-SVs to the AZ membrane decreased upon stimulation (Fig. 7A), indicating recruitment of SVs to the AZ membrane. Moreover, SVs were found closer to the PD upon light stimulation (Fig. 7B), likely bringing them close to the voltage-gated Ca^2+^ channels that are situated underneath to the PD (Neef et al., 2018; Pangrsic et al., 2018; Wong et al., 2014). This trend was only significant for ChR2 LongStim (all values can be found in Table 3). We conclude that upon stimulation, SVs are recruited more tightly to the AZ membrane and potentially closer to the Ca^2+^ channels. Similar results were obtained for the individual genotypes (Fig. 7-figure supplement 1).

**Figure 7.**
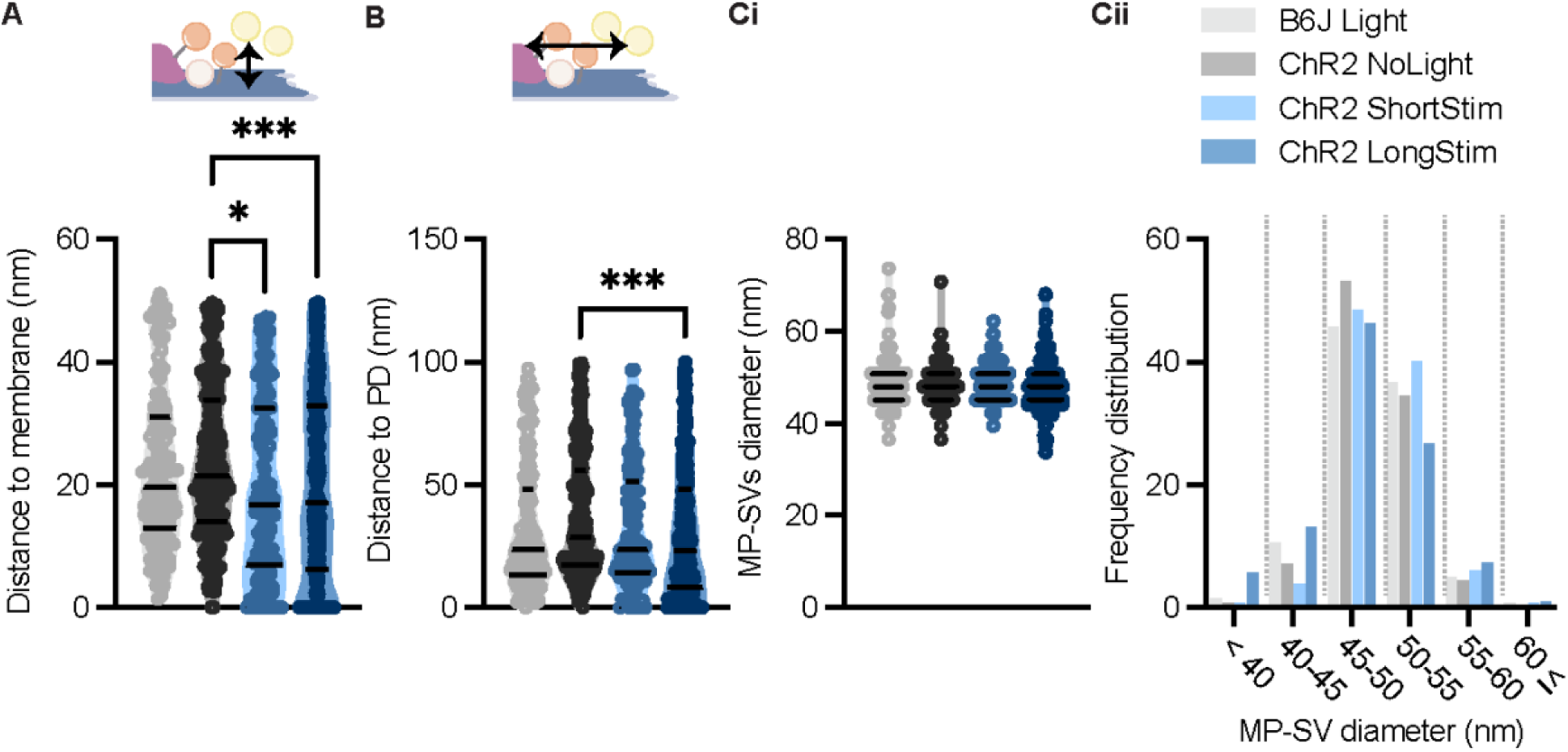
MP-SVs come closer to the AZ membrane and the presynaptic density upon light stimulation. **(A)** MP-SVs distance to the AZ membrane. **(B)** MP-SVs distance to the PD. **(Ci)** Diameter of MP- SVs quantified from the outer rim to the outer rim. **(Cii)** Frequency distribution of SV diameter of all MP-SVs. Data are presented in mean ± SEM. *p < 0.05, **p < 0.01, ***p < 0.001 and ****p < 0.0001. Statistical test: one-way ANOVA followed by Tukey’s test (parametric data) and KW test followed by Dunn’s test (non-parametric data). B6J Light: *n* = 15 ribbons, *N_animals_* = 2; ChR2 NoLight: *n =* 17 ribbons *N_animals_* = 4; ChR2 ShortStim: *n* = 11 ribbons, *N_animals_* = 1; ChR2 LongStim: *n =* 26 ribbons, *N_animals_* = 4.

### SV diameters remain largely unchanged in the MP-SV pool

In order to approach possible release mechanisms for IHC ribbon synapses, we determined the SV diameter for all MP-SVs. Previous studies using 15 min K^+^ stimulation already excluded a large increase of SV sizes upon prolonged stimulation close to the AZ membrane (Chakrabarti et al., 2018; Chapochnikov et al., 2014). However, early exocytosis phases could not be monitored at IHC ribbon synapses up to now. If homotypic or compound fusion takes place, one would expect an increase in diameter of SV close to the AZ membrane (He et al., 2009; Lenzi et al., 2002). We found no differences in the diameters of the MP-SVs between the stimulated and non-stimulated conditions (Fig. 7C; values and details for statistics in Table 3) We investigated the SV diameter distribution in more detail by sorting all MP-SVs into different bins. We examined small SVs > 40 nm in diameter as well as large SVs ≤ 60 nm, and the frequency distribution between 45 and 60 in 5 nm steps. There were no obvious shifts in the frequency distributions of the SV diameters (Fig. 7Cii). In conclusion, homotypic SV fusion events do not seem to take place among MP-SVs of the IHC synapse under our stimulation paradigms.

## Discussion

In the current study, we established Opto-HPF with a millisecond range physiological stimulation, followed by FS and ET for structure-function analysis of IHC ribbon synapses. This enabled near-to-native state preservation of the ultrastructure of exocytic steps occurring within milliseconds and offered a closer correlation to cell-physiological stimulation paradigms widely used in the field. Patch-clamp recordings validated the photoresponses of ChR2-expressing IHCs and demonstrated optogenetically triggered glutamate release. Further, we provide a strategy for precise synchronization of HPF with optogenetic stimulation. In summary (Fig. 8), our analysis revealed a stimulation-dependent accumulation of docked SVs at IHC AZs. Moreover, we found a slight reduction of the distance of non-docked SVs to the AZ membrane and the PD, even more prominent with longer stimulation duration. Finally, with this physiological stimulation, we did not observe large SVs or other morphological correlates of potential homotypic fusion events in the MP-SV pool.

**Figure 8.**
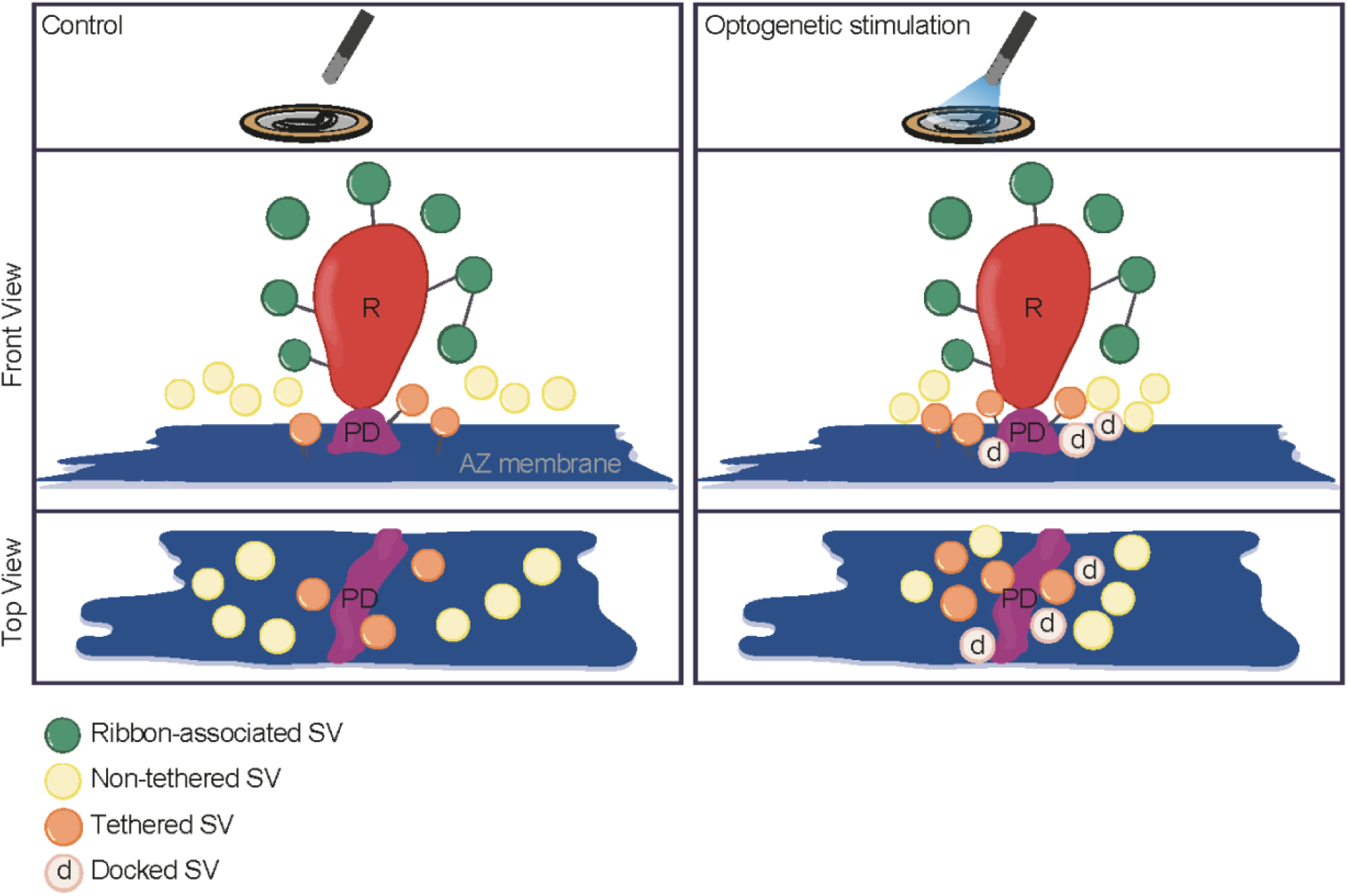
Summary. Optogenetic stimulation of IHCs mobilize SVs more tightly to the AZ membrane and potentially closer to the Ca^2+^ channels. The proportion of docked and tethered SVs increased upon stimulation duration, while the total count of MP-SVs and RA-SVs stayed stable. The distance of MP-SVs to the AZ membrane decreased with stimulation duration.

### Validation of Opto-HPF in IHCs

Using immunofluorescence microscopy, immunogold electron microscopy and patch-clamp recordings, we observed an efficient and functional expression of ChR2 (H134R) in IHCs of mouse lines that employed Vglut3-dependent Cre expression. Blue light pulses evoked photocurrents and -depolarizations similar to previous reports in other cell types (Boyden et al., 2005; Cardin et al., 2010; Hernandez et al., 2014; Kittelmann et al., 2013; Nikolic et al., 2009). Our patch-clamp recordings of ChR2-expressing IHCs or of the postsynaptic bouton revealed that a 10 ms light pulse of 6-16 mW/mm^2^ was sufficient to i) depolarize the IHC by 50 mV within 10 to 20 ms and ii) trigger EPSCs at individual synapses with latencies of 15-20 ms. Longer stimulations of similar irradiance decreased the time to peak of photodepolarization and EPSC latencies and resulted in a sustained release. Part of this sustained release is attributed to the slow repolarization of the IHCs (>40 ms) due to the presence of K^+^ channel blockers (TEA-CL and Cs^+^ in the present study). Based on the charge of the light-evoked EPSCs, the recorded EPSCs most likely reflect the fusion of several SVs at the individual AZ. Under the premise that an individual SV leads to a charge transfer ranging from 50 to 600 fC (Huang and Moser, 2018; Rutherford et al., 2012), our optogenetic stimulation of IHCs triggers the release of more than 10 SVs on the recorded AZs. This number of released SVs and the presence of an immediate plateau indicates depletion of the RRP even after short and mild light stimulations.

In order to arrive at a reliable Opto-HPF operation when using the HPM100, we added further functionalities to the machine. The pneumatic pressure sensor, which was placed in close proximity to the freezing chamber, accurately enabled us to calculate the time point when pressurized LN_2_ entered the freezing chamber. This allowed a correlation to the data of internal pressure and temperature sensors, whereas the other sensors provided less reliable signals. To achieve short and long light stimulations in the HPM, we chose a single light pulse with different onset time points. According to the sensor curves, we obtained light stimulations between 17 and 76 ms. Our ShortStim (∼20 ms) and LongStim (∼50 ms) Opto-HPF paradigms aimed to capture ultrastructural correlates of such phasic and sustained exocytosis and matched stimulus durations widely used in electrophysiology of hair cell exocytosis (e.g. Cho et al., 2011; Goutman and Glowatzki, 2007; Johnson et al., 2017; Michalski et al., 2017; Moser and Beutner, 2000; Parsons et al., 1994; Schnee et al., 2005). IHC patch-clamp indicates that RRP is released within the first 20 ms of step depolarization (Goutman and Glowatzki, 2007; Moser and Beutner, 2000) while longer depolarizations recruit additional SVs for sustained exocytosis (Goutman and Glowatzki, 2007; Moser and Beutner, 2000). Furthermore, instead of applying trains of short light pulses as typically used for neuronal cell types (Berndt et al., 2011; Boyden et al., 2005; Imig et al., 2020; Ishizuka et al., 2006; Kleinlogel et al., 2011; Lin et al., 2009), we opted for a continuous light pulse to mimic a step-like receptor potential of IHCs in the high frequency cochlea of the mouse (Russell and Sellick, 1978). The ultrastructural findings upon light stimulation in the HPM undoubtedly reflect snapshots of exocytosis at the IHC synapse.

### Resolving IHCs synaptic vesicle pools with Opto-HPF

Significant efforts have been made to address the mechanisms of SV release at different ribbon synapses by studying morphologically defined SV populations using ET. These studies proposed that the SVs situated close to the AZ membrane represent the ‘’ultrafast release pool’’ (Lenzi and von Gersdorff, 2001; Lenzi et al., 1999), and SVs further away around the ribbon are accessible for slower release (Lenzi et al., 1999).

Capacitance measurements (Beutner and Moser, 2001; Johnson et al., 2005; Khimich et al., 2005; Moser and Beutner, 2000; Pangrsic et al., 2010), fluorescence imaging (Griesinger et al., 2005; Özçete and Moser, 2020) and recordings from single spiral ganglion neurons (Buran et al., 2010; Frank et al., 2010; Goutman and Glowatzki, 2007; Jean et al., 2018; Jung et al., 2015a; Peterson et al., 2014) propose an RRP with a size of between 4 to 45 vesicles per AZ which partially depletes with a time constant of 3 to 54 ms. The broad range of size and release kinetics estimates results from differences in methods, stimulus paradigms and experimental conditions as well as in assumptions of model-based data analysis. Moreover, heterogeneity of AZs may also play a role. These physiological estimates of RRP size enclose the number of approximately 10 MP-SVs. Yet, docking of SVs, often considered to be the ultrastructural correlate of fusion competence, is virtually absent from IHC AZs in non-stimulated conditions (present study and (Chakrabarti et al., 2018)). Moreover, in contrast to the physiological evidence for a partial RRP depletion, IHC ribbon synapses did not display a significant reduction in MP-SVs upon optogenetic stimulation on the ultrastructural level. Strikingly, instead and contrary to conventional and retinal ribbon synapses (Borges-Merjane et al., 2020; Imig et al., 2020; Watanabe et al., 2013b), there is a prominent increase in docked SVs (present study and (Chakrabarti et al., 2018)).

Finding an accumulation of docked SVs upon strong depolarization seems puzzling given estimated rates of SV replenishment and subsequent fusion of 180-2000 SV/s at IHC ribbon synapses (Buran et al., 2010; Goutman and Glowatzki, 2007; Jean et al., 2018; Pangrsic et al., 2010; Peterson et al., 2014; Schnee et al., 2011; Strenzke et al., 2016). Indeed, such high speed and indefatigable SV release enable firing up to approximately 100 spikes/s in the quiet and steady state firing of up a few hundred spikes/s upon strong sound stimulation (Buran et al., 2010; Evans, 1972; Huet et al., 2016; Jean et al., 2018; Kiang et al., 1965; Liberman and Kiang, 1978; Schmiedt, 1989; Taberner and Liberman, 2005). Do these docked SVs represent release ready SVs that are more likely detected upon massive turnover? Do they reflect “kiss and stay” release events or limited clearance of vesicles following release? Does the lack of docked SVs at resting IHC synapses reflect a rapid undocking process?

SV clearance of the AZ (Neher and Sakaba, 2008) has been suggested as a potentially rate-limiting mechanism of sustained exocytosis in IHCs of mice with mutations in the genes coding for otoferlin (Chakrabarti et al., 2018; Pangrsic et al., 2010; Strenzke et al., 2016) or endocytic proteins (Jung et al., 2015b; Kroll et al., 2019; Kroll et al., 2020). While our EPSC recordings suggest the ongoing release of neurotransmitter beyond 20 and 50 ms after light onset, limited clearance of the release sites cannot be excluded. The concept implies full collapse fusion followed by clearance of SV proteolipid from the release site for it to engage a new coming SV. Indeed, full collapse fusion followed by clathrin- and dynamin-dependent endocytosis has been indicated for IHCs (Grabner and Moser, 2018; Neef et al., 2014; Tertrais et al., 2019). Yet, unlike for retinal ribbon synapses (Zampighi et al., 2006; Zampighi et al., 2011) we did not observe omega- profiles or hemifusion states of SVs at IHCs AZ opposing the postsynaptic density.

While we have favored the hypothesis that, eventually, fusion pore initiated release typically proceeds to full collapse fusion (Chapochnikov et al., 2014), there is support for “kiss and run” exocytosis (Alabi and Tsien, 2013) to occur at IHCs from reports of ultrafast endocytosis (time constant ∼300 ms) (Beutner et al., 2001; Neef et al., 2014; Tertrais et al., 2019) and cell-attached capacitance measurements (Grabner and Moser, 2018). Could the accumulation of docked SVs during stimulation then represent “kiss and run” or “kiss and stay” (Shin et al., 2018) release events? Unfortunately, cell-attached membrane capacitance recordings from IHCs did not resolve fusion pores (Grabner and Moser, 2018), probably owing to the small SV capacitance (40 aF). Future work including super-resolution imaging (Shin et al., 2018) and/or Opto-HPF on IHCs with genetic or pharmacological interference might shed light on the existence of a prevalence of “kiss and run” or “kiss and stay” at IHC synapses. Freezing times between 5 to 15 ms after the light onset seem necessary to further address this hypothesis.

Finally, a recent study using electrical stimulation and HPF in hippocampal neurons reported full replenishment of docked SVs within 14 ms, but this docking state was only transient, and SVs could potentially undock during the next 100 ms (Kusick et al., 2020). Indeed, physiological evidence for reversible priming and docking has been reported for various neurosecretory preparations (e.g. Dinkelacker et al., 2000; He et al., 2017; Nagy et al., 2004; Smith et al., 1998). The balance of Ca^2+^ and otoferlin-dependent replenishment of docked and primed SVs with SV fusion and/or undocking/depriming would then set the “standing RRP” (Pangrsic et al., 2010) and the abundance of docked SVs. Besides increased docking, the decreased distance between SVs and the plasma membrane and presynaptic density upon light stimulation supports the previously proposed sequence of events at IHC ribbon synapses (Chakrabarti et al., 2018). The sequence for SV release involves tethering and subsequent docking, similarly to originally described in conventional synapses using cryo-ET (Fernández-Busnadiego et al., 2013). Aside from RIMs, which support SV tethering to the AZ membrane at conventional (Fernández-Busnadiego et al., 2013) and IHC ribbon synapses (Jung et al., 2015a), otoferlin (Pangrsic et al., 2010; Vogl et al., 2015) rather than neuronal SNAREs (Nouvian et al., 2011; but see Safieddine and Wenthold, 1999) and members of the Munc-13/CAPS families of priming factors (Vogl et al., 2015) seem to contribute to preparing SVs for fusion. It will be interesting for future studies to further decipher the underlying molecular machinery at the hair cell synapses and to test whether and to what extent tethering and docking are reversible processes.

## Conclusion

Significant efforts have been made to address the release scenarios at ribbon synapses by EM (Chakrabarti et al., 2018; Chapochnikov et al., 2014; Lenzi et al., 2002; Matthews and Sterling, 2008; von Gersdorff et al., 1996; Zampighi et al., 2011). This study offers an experimental approach for structure-function correlation at IHC ribbon synapses. We conclude that activation of ChR-2 rapidly depolarizes IHCs and triggers release within few milliseconds in response to brief light flashes. This constitutes a non-invasive approach that overcomes the low temporal resolution of the conventional high K^+^ depolarization used for electron microscopy of IHC synapses thus far. Combining optogenetic stimulation with high-pressure freezing appears as a promising technique to achieve the temporal and spatial resolution required to study the short-term cellular processes occurring during exocytosis and endocytosis. Notably, we did not observe events that resemble homotypic SV fusion or cumulative fusion close to the AZ membrane of the synaptic ribbon. Further, the RA- as well as the MP-SV pools stayed stable or were rapidly replenished, rather a decrease of the fraction of non-tethered SVs within the MP-SV pool was observed. Finally, the absence of docked SV in non-stimulated IHCs might speak for uniquantal release, possibly also due to fast undocking, to prevail for spontaneous release events.

## Acknowledgements

We thank A.J. Goldak, S. Langer, S. Gerke, and C. Senger-Freitag for expert technical assistance. We would like to thank P. Wenig for the technical support in establishing the HPM setup. We would like to thank T. Mager for helpful discussion on the irradiance and the company Leica for support with the sensors.

## Author contributions

RC established and performed Opto-HPF and analysis of the ultrastructural data, electron tomography (ET), pre-embedding immunogold labeling and contributed to the immunohistochemistry. LMJT performed all cell physiology and according analysis and contributed to the immunohistochemistry and prepared figures. LS performed Opto-HPF, analysis of the ultrastructural data, ET and prepared figures. MRC performed immunofluorescence, analysis of ribbon numbers, helped in pre-embedding immunogold analysis and Opto-setup and prepared figures. GH programmed the MATLAB GUI, MATLAB interface, installed the sensors to the Opto-HPF. MS performed part of the ultrastructural analysis and ET, and EF performed part of the ET analysis. KB established and performed the irradiance measurement. ÖDÖ designed primers for genotyping. TP contributed to the KI line. SM supported HPF and statistical analysis of the SV diameters. JN supervised LMJT cell physiology, contributed to immunostainings. MG performed the statistical analysis for the SV diameters. FO developed nanogold coupled nanobodies and helped to design the labeling protocol. TM designed the study and supervised LMJT/ cell physiology. CW designed the study and supervised Opto-HPF and immunostainings and prepared figures. CW, TM and LMJT wrote the manuscript with the help of all authors.

## Additional Files

Source Code 1: IMARIS custom plug-ins for the analysis of Figure 1D

---

**Figure 1-figure supplement 1.**
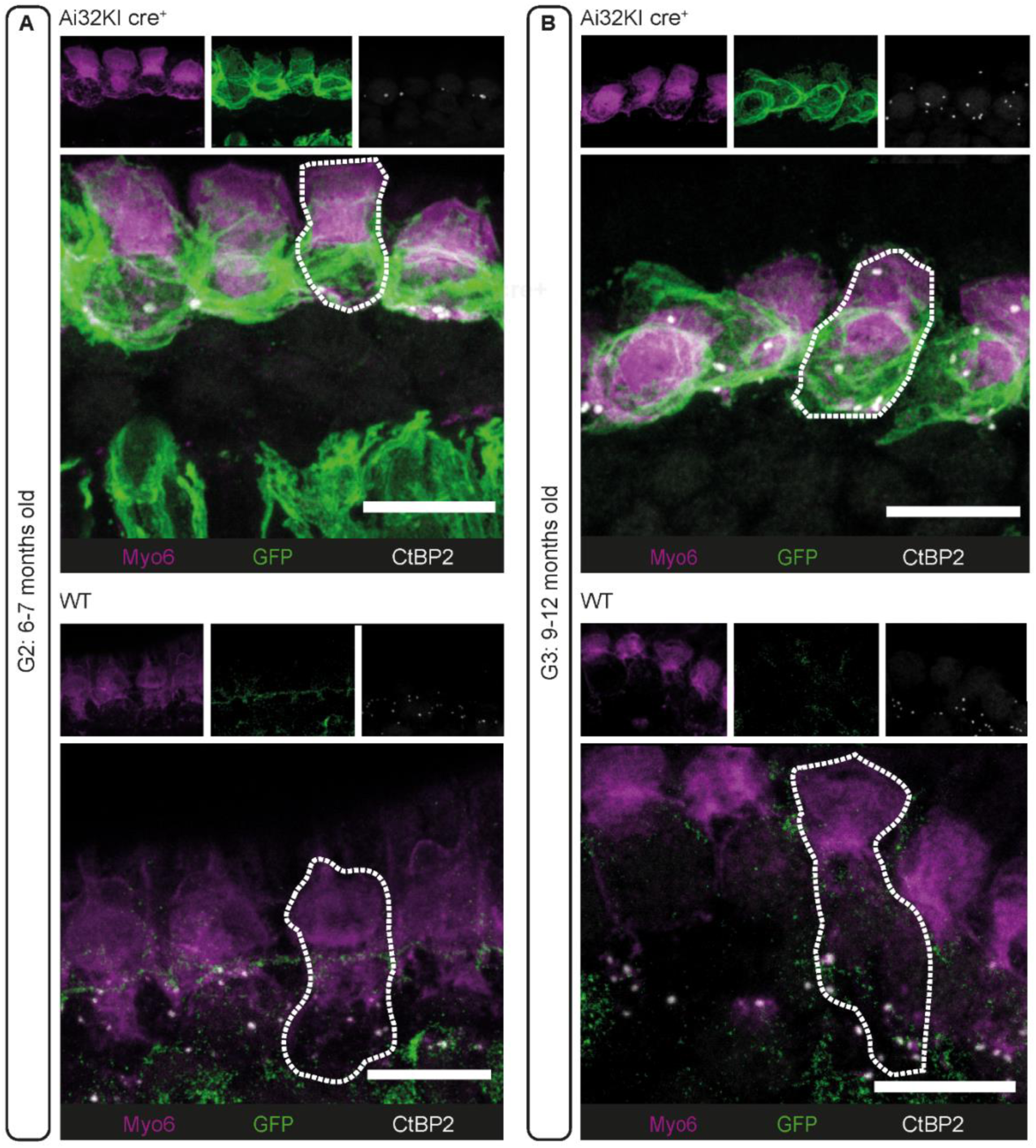
Long term expression of ChR2 at IHC plasma membrane. Maximal projection of confocal *z*-stacks of **(A)** G2 (6-7 months-old mice) and **(B)** G3 (9-12 months- old mice) IHCs from the apical turn of organs of Corti from ChR2-expressing mice (*Ai32KI cre^+^*; upper panels) and their respective controls (*WT*; lower panels). Some IHCs are outlined with a dotted line. ChR2 expression at the membrane is visualized by GPF labeling (green). Myo6 (magenta) immunostaining was used to delineate the IHC and Ctbp2 to visualize the ribbons (white). Ectopic GFP expression (**A**, upper panel) in spiral ganglion fibers is indicative of unspecific *cre* recombination (see Materials & Methods). Scale bars, 10 µm.

**Figure 2-figure supplement 1.**
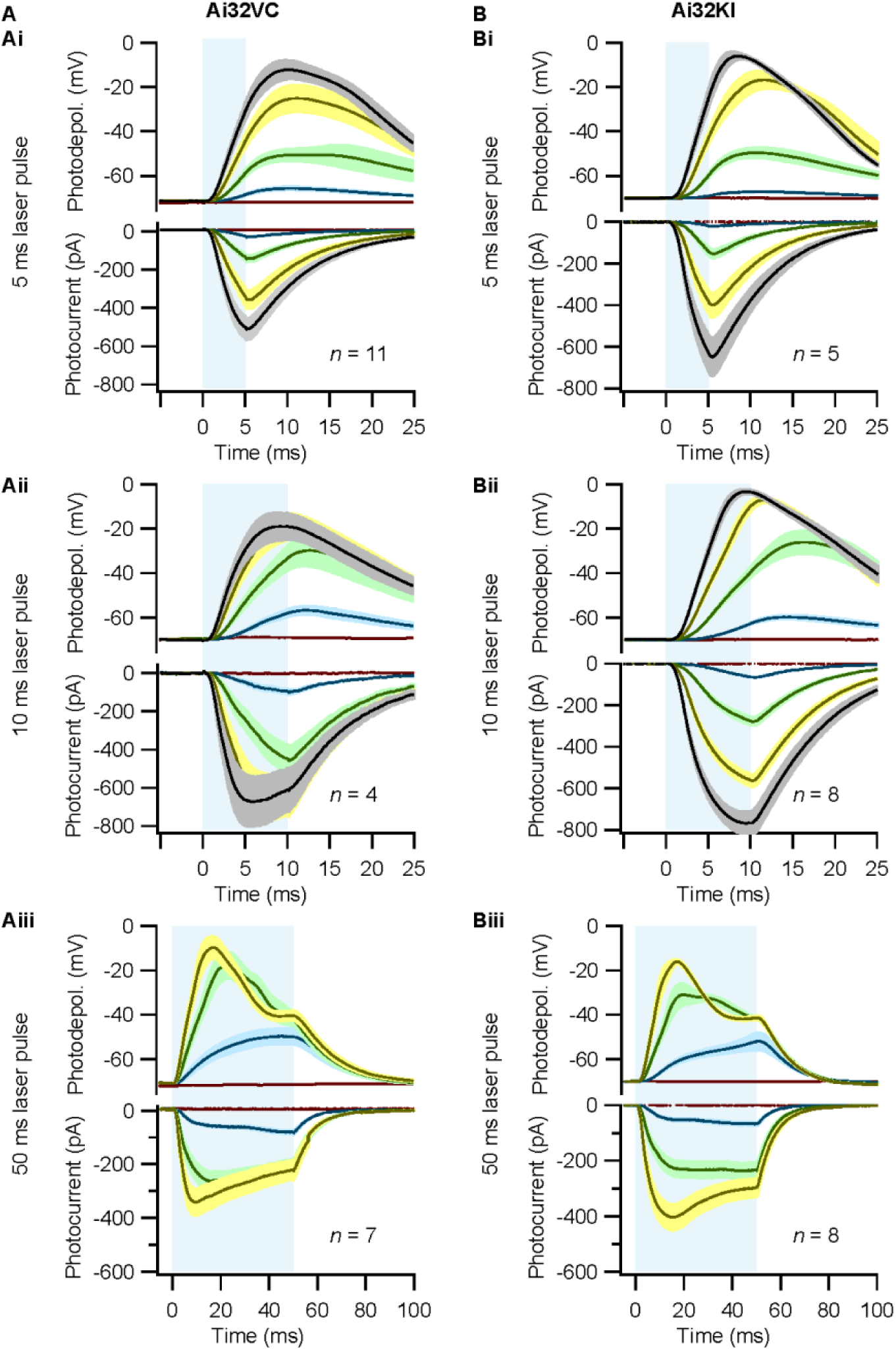
Comparison of optogenetic stimulation of IHCs from *Ai32VC cre^+^* and *Ai32KI cre^+^* mice. IHCs expressing ChR2 (*Ai32VC cre^+^* left and *Ai32KI cre^+^* right) were optogenetically stimulated by 473 nm light pulses of increasing irradiance (mW/mm^2^). **(A-C)** Average photocurrents (lower panel) and photodepolarizations (upper panel) of patch-clamped IHCs during 5 ms (A), 10 ms (B) and 50 ms light pulses of increasing irradiances (color coded as in Fig. 2). Mean is displayed by the continuous line and ± SEM by the shaded area. For 5 ms: *n_Ai32VC_* = 11, *N_animals Ai32VC_* = 6; *n_Ai32KI_* = 5, *N_animals Ai32KI_* = 1. For 10 ms: *n_Ai32VC_* = 4, *N_animals Ai32VC_* = 1; *n_Ai32KI_* = 8, *N_animals Ai32KI_* = 3. For 50 ms: *n_Ai32VC_* = 7, *N_animals Ai32VC_* = 3; *n_Ai32KI_* = 8, *N_animals Ai32KI_* = 3.

**Figure 4-figure supplement 1.**
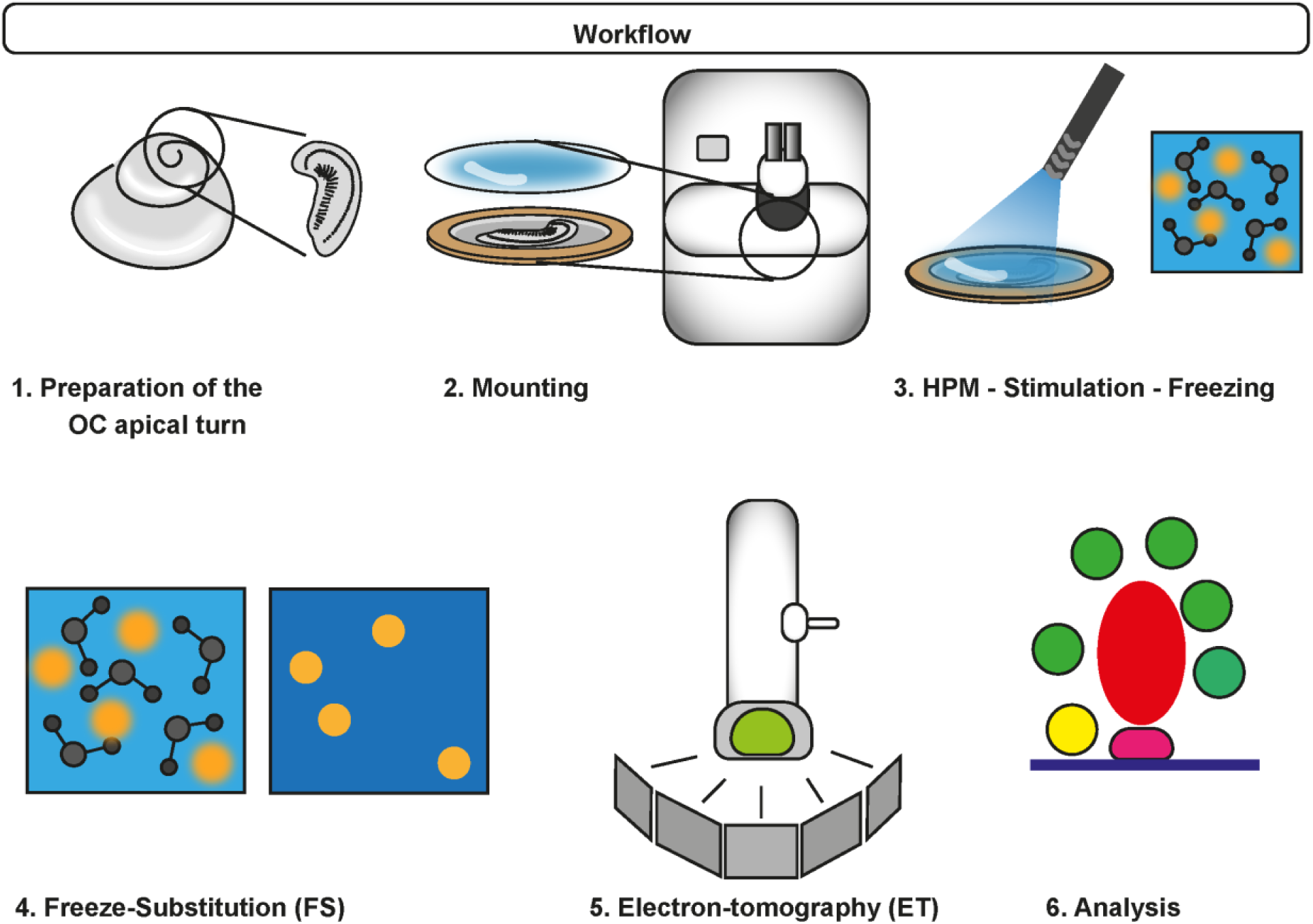
Workflow of Opto-HPF. After dissection of the organ of Corti (OC), the sample is mounted and inserted in the HPM. The blue light stimulation occurs in the HPM freezing chamber with subsequent freezing. Finally, freeze-substitution is performed followed by ET and data analysis.

**Figure 6-figure supplement 1.**
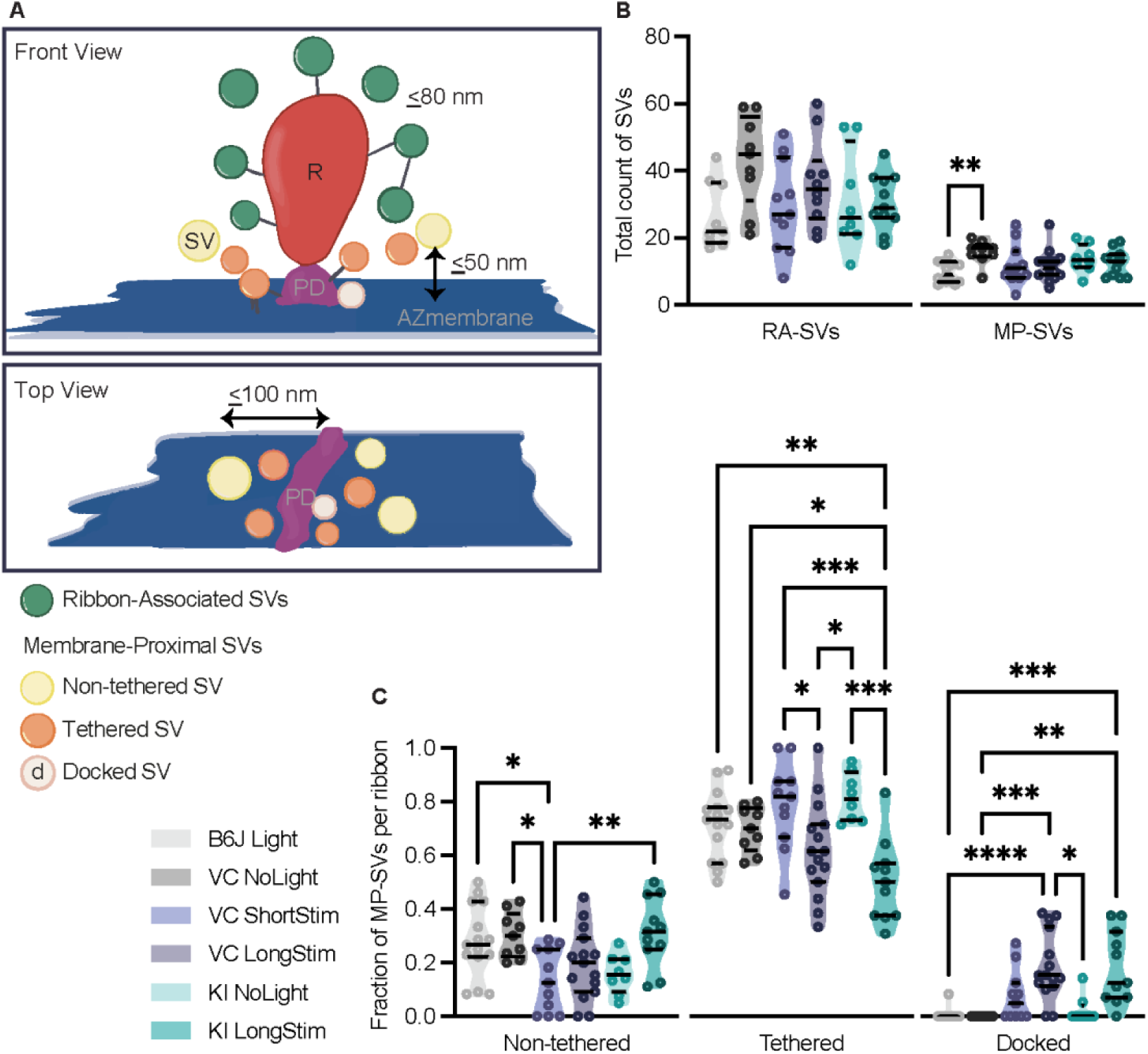
Analysis of morphologically defined vesicle pools for each genotype. **(A)** Schematic illustration of a ribbon synapse (not drawn to scale) showing the parameters taken into account for the analysis of the different vesicle pools. Membrane-proximal (MP)-SVs constitute the first-row of vesicles within 50 nm membrane-to-membrane distance from the AZ- membrane (blue) and 100 nm from the presynaptic density (PD, pink). Non-tethered SVs are in yellow, tethered in orange and docked in light pink. For Ribbon-associated (RA)-SVs, vesicles (green) within 80 nm from the ribbon (R, in red) are included. **(B)** Total count of SVs per pool (RA- and MP-SV pools), per ribbon. **(C)** Fraction of non-tethered, tethered and docked MP-SVs per ribbon for the controls as well as for Ai32VC_ShortStim, Ai32VC_LongStim and Ai32KI_LongStim Data are presented in mean ± SEM. *p < 0.05, **p < 0.01, ***p < 0.001 and ****p < 0.0001. Statistical test: one-way ANOVA followed by Tukey’s test (parametric data) and KW test followed by Dunn’s test (non-parametric data). MP-SV pool: B6J_LongStim: *n* = 15 ribbons, *N_animals_* = 2; Ai32VC_NoLight: *n =* 9 ribbons *N_animals_* = 2; Ai32VC_ShortStim: *n* = 11 ribbons, *N_animals_* = 1. Ai32VC_LongStim: *n =* 15 ribbons, *N_animals_* = 2. Ai32KI_LongStim: *n =* 11 ribbons *N_animals_* = 2; Ai32KI_NoLight: *n =* 8 ribbons *N_animals_* = 2. RA-SV pool: B6J_LongStim: *n* = 9 ribbons, *N_animals_* = 1; Ai32VC_NoLight: *n =* 9 ribbons *N_animals_*= 2; Ai32VC_ShortStim: *n* = 11 ribbons, *N_animals_* = 1; Ai32VC_LongStim: *n =* 10 ribbons, *N_animals_* = 1; Ai32KI_LongStim: *n =* 11 ribbons *N_animals_* = 2; Ai32KI_NoLight: *n =* 8 ribbons *N_animals_*= 2.

**Figure 7-figure supplement 1.**
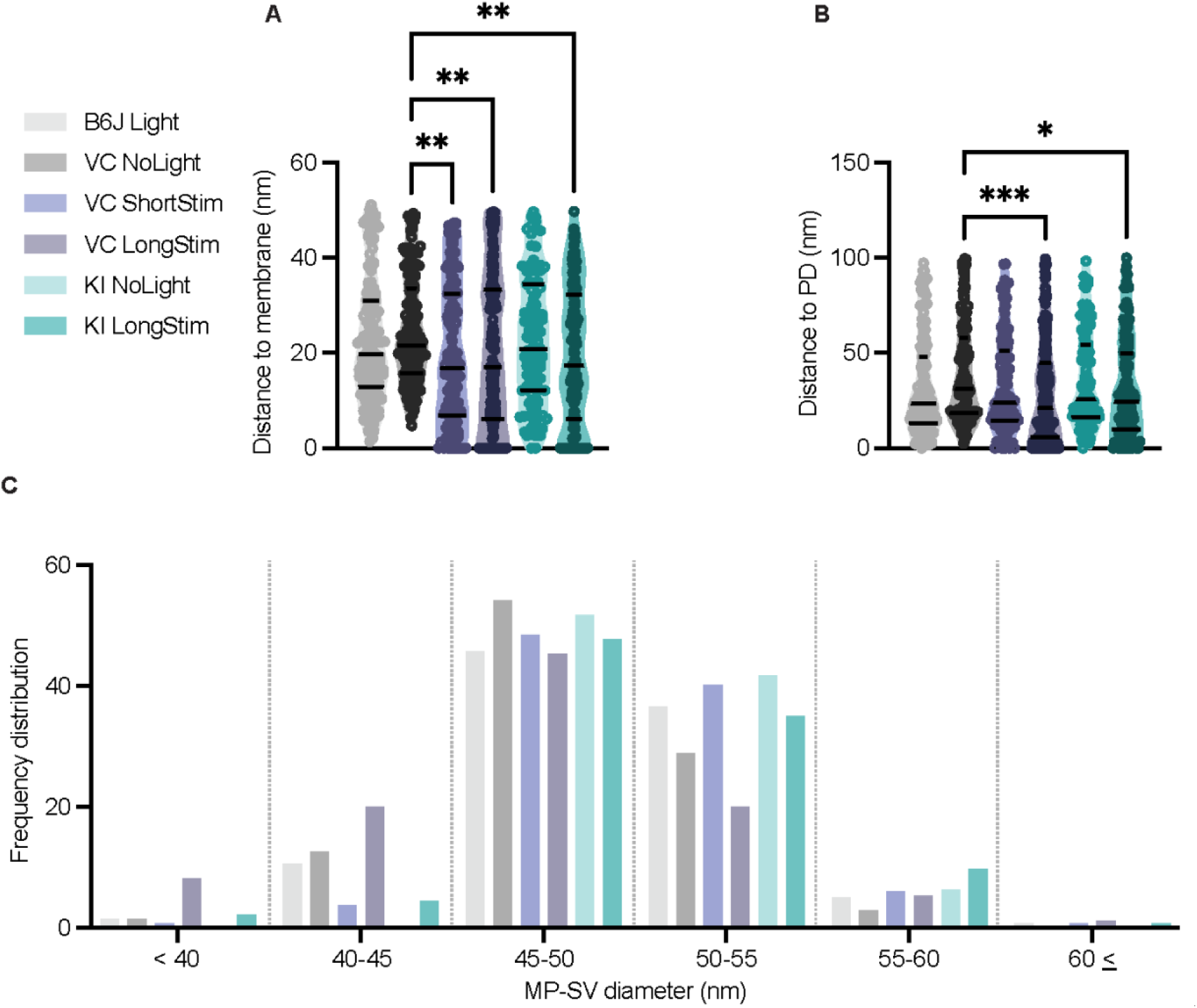
Distances of MP-SVs to the AZ membrane and the presynaptic density as well as their diameters. **(A)** MP-SVs distance to the AZ membrane. **(D)** MP-SVs distance to the PD. **(E)** Frequency distribution of SV diameter of all MP-SVs. Data are presented in mean ± SEM. *p < 0.05, **p < 0.01, ***p < 0.001 and ****p < 0.0001. Statistical test: one-way ANOVA followed by Tukey’s test (parametric data) and KW test followed by Dunn’s test (non-parametric data). B6J_LongStim: *n* = 15 ribbons, *N_animals_* = 2; Ai32VC_NoLight: *n =* 9 ribbons *N_animals_* = 2; Ai32VC_ShortStim: *n* = 11 ribbons, *N_animals_* = 1; Ai32VC_LongStim: *n =* 15 ribbons, *N_animals_* = 2; Ai32KI_ NoLight: *n* = 8 ribbons, *N_animals_* = 2; Ai32KI_ LongStim: *n* = 11 ribbons, *N_animals_* = 2

